# Microbial association networks give relevant insights into plant pathobiomes

**DOI:** 10.1101/2020.02.21.958033

**Authors:** Charlie Pauvert, Tania Fort, Agnès Calonnec, Julie Faivre d’Arcier, Emilie Chancerel, Marie Massot, Julien Chiquet, Stéphane Robin, David A. Bohan, Jessica Vallance, Corinne Vacher

## Abstract

Interactions between plant pathogens and other plant-associated microorganisms regulate disease. Deciphering the networks formed by these interactions, termed pathobiomes, is crucial to disease management. Our aim was to investigate whether microbial association networks inferred from metabarcoding data give relevant insights into pathobiomes, by testing whether inferred associations contain signals of ecological interactions. We used Poisson Lognormal Models to construct microbial association networks from metabarcoding data and then investigated whether some of these associations corresponded to interactions measurable in co-cultures or known in the literature, by using grapevine (*Vitis vinifera*) and the fungal pathogen causing powdery mildew (*Erysiphe necator*) as a model system. Our model suggested that the pathogen species was associated with 23 other fungal species, forming its putative pathobiome. These associations were not known as interactions in the literature, but one of them was confirmed by our co-culture experiments. The yeast *Buckleyzyma aurantiaca* impeded pathogen growth and reproduction, in line with the negative association found in the microbial network. Co-cultures also supported another association involving two yeast species. Together, these findings indicate that microbial networks can provide plausible hypotheses of ecological interactions that could be used to develop microbiome-based strategies for crop protection.

## INTRODUCTION

Plant growth and health depend on their associations with a large number of microorganisms that interact with each other (Vandenkoornhuyse et al. 2015; Hassani et al. 2018). Among all these microorganisms, those that interact with pathogens and regulate diseases are particularly important and were recently termed the plant’s pathobiome (Vayssier-Taussat et al. 2014; Brader et al. 2017; Bass et al. 2019). Some pathobiome members form a barrier that limits pathogen development through direct antagonistic interactions (Arnold et al. 2003; Kemen 2014; Durán et al. 2018; Li et al. 2019), while others can prime the plant immune system (Vogel et al. 2016; Lee et al. 2017; Hacquard et al. 2017). In turn, a successful pathogen invasion can disrupt the plant microbiota and drive higher community heterogeneity (Zaneveld et al. 2017). This increase in heterogeneity among infected microbiotas is termed the Anna Karenina principle, based on the first sentence of Leo Tolstoy’s book: “Happy families are all alike; every unhappy family is unhappy in its own way”. This can be transposed to microbiology as: “All healthy microbiomes are similar; each dysbiotic microbiome is dysbiotic in its own way” (Zaneveld et al. 2017).

Deciphering microbial interactions within pathobiomes, and determining what environmental factors shape those interactions, will be an important step towards improved plant health protection. A better understanding of plant pathobiomes will allow us to reduce our reliance on chemical pesticides, through the discovery of novel biocontrol agents (Poudel et al. 2016) and cultural practices fostering the protective microbiota (Hartman et al. 2018). To achieve this aim, research is needed at the interface between plant pathology, microbial community ecology, culturomics, metagenomics and big data (Vannier et al. 2019). Culturable and unculturable microorganisms associated with healthy and diseased plants can be described using metabarcoding approaches (Abdelfattah et al. 2018; Nilsson et al. 2019). Network inference methods can then be applied to reconstruct microbial association networks from metabarcoding data (Faust and Raes 2012; Biswas et al. 2016; Vacher et al. 2016; Layeghifard et al. 2017). These networks, in which nodes correspond to microbial taxa and links to direct statistical associations between their sequence counts, constitute hypotheses of microbial interactions (Jakuschkin et al. 2016; Poudel et al. 2016). Finally, microbiological experiments could then be used to test these interaction hypotheses (*e.g.* Lima-Mendez et al. 2015; Biswas et al. 2016; Wang et al. 2017; Das et al. 2018; Durán et al. 2018; Gao et al. 2018) and develop microbiome-based plant protection strategies.

Despite its potential interest to agriculture and environment, the application of network inference methods to pathobiome research is still in its infancy. Several methodological issues must be overcome to generate robust and relevant hypotheses of microbial interactions from metabarcoding data because statistical associations between sequence counts do not directly reflect ecological interactions (*e.g.* competition, parasitism) between microorganisms (Weiss et al. 2016; Derocles et al. 2018; Röttgers and Faust 2018). For instance, the compositional nature of metabarcoding data induces statistical associations between sequence counts that are not related to any ecological process (Friedman and Alm 2012; Gloor et al. 2017). Environmental filtering may also generate statistical associations between microbial taxa abundances that are not triggered by ecological interactions but environmental variations (Berry and Widder 2014; Vacher et al. 2016; Derocles et al. 2018; Röttgers and Faust 2018). Several recent methods of network inference deal explicitly with these two issues, including HMSC (Hierarchical Modelling of Species Communities; Ovaskainen et al. 2017) and PLN (Poisson LogNormal Model; Chiquet et al. 2018; 2019). However, their relevance to pathobiome research has yet to be demonstrated.

The aim of this study was to deepen our knowledge of the dynamics of species and interactions within plant pathobiomes. We tested the following hypotheses: (H1) successful infection events destabilize plant-associated microbial communities and increase their heterogeneity (Anna Karenina principle); (H2) interactions among microorganisms within pathobiomes can be detected by inferring microbial networks from metabarcoding data and environmental covariates; and, (H3) cropping system influences the abundance of microorganisms forming pathobiomes. These hypotheses were tested using grapevine, *Vitis vinifera*, and the fungal obligate biotroph pathogen causing powdery mildew, *Erysiphe necator* (Armijo et al. 2016, Gadoury et al. 2012), as a model system. To maximize the diversity of grapevine-associated microbial communities and thus the possibility of discovering microorganisms interacting with the pathogen, we conducted the study in an untreated vineyard plot (Figure 1). We inoculated the pathogen species on several plants at the beginning of the growing season and then collected sporulating and visually healthy leaves on three dates (40, 62 and 77 days post-inoculation). Illumina MiSeq sequencing was used to characterize fungal communities in three tissue types: visually healthy leaf blade of non-sporulating leaves (HNI); visually healthy leaf blades of sporulating leaves (HI); and, disease spots (DI). We used the PLN model (Chiquet et al. 2018; 2019) to construct a network of associations between foliar fungal species from the metabarcoding dataset, and then used a combination of co-cultures and text-mining to test whether these statistical associations represented ecological interactions. The ecology of pathobiome species was characterized by searching for their presence in bark, ground cover and upper soil, and by assessing their response to cover cropping (CC) and weed removal (NCC; Figure 1).

**Figure 1.**
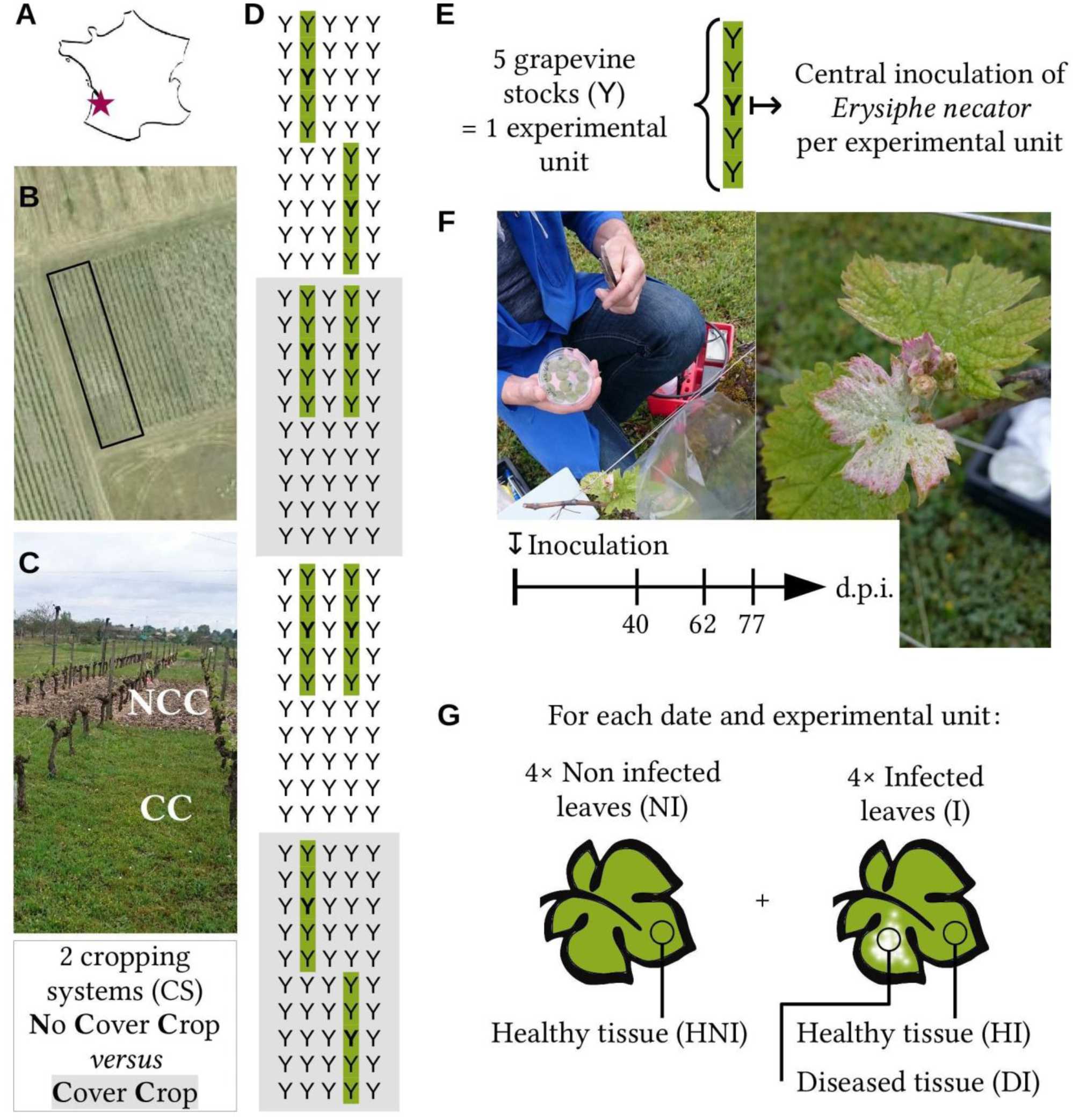
**Experimental design.** The study took place (A) in an experimental vineyard located near Bordeaux, France, (B) in a sub-plot of 5 untreated vine rows, (C) with two cropping systems differing in the presence or not of cover crop (CC *versus* NCC) in the inter-rows. (D) Sampling occurred in eight experimental units each composed of five adjacent vines (underlined in green). (E) The causal agent of grape powdery mildew, *Erysiphe necato*r, was inoculated on the central vine of each experimental unit at the beginning of the vegetative season. (F) We performed three sampling campaigns on June 1, 23 and July 5 (corresponding to 40, 62 and 77 days post inoculation (dpi), respectively). (G) For each campaign, leaves with and without visible symptoms were collected to analyze healthy foliar tissues and foliar tissue from diseased spots.

## RESULTS

### Pathogen abundance and fungal load tend to be lower in visually healthy tissues

The grapevine foliar fungal community was composed of 4148 fungal amplicon sequence variants (ASVs; Callahan et al. 2017), representing 10195266 sequences. 1454 of these ASVs, representing 71.4% of the sequences, were assigned at the species level using the UNITE database (UNITE Community 2019). These ASVs corresponded to 547 fungal species, of which 306 were Ascomycetes and 241 Basidiomycetes. The pathogen *E. necator* was among the three most abundant fungal species, even in visually healthy tissues. It ranked first in sporulating leaves, in both visually healthy tissues and disease spots. It ranked third in leaves with no visible symptom, after *Mycosphaerella tassiana* and *Filobasidium wieringae* (Table 1). The relative abundance of *E. necator* varied significantly through the vegetative season (Table 2A), although we collected leaves of the same age on the three sampling dates. On the first and second sampling dates (both in June), the pathogen represented 39% and 39.6% of all fungal sequences, respectively. As expected, it was significantly more abundant in disease spots than in visually healthy tissues (Table 2A and Figure S1A). The total fungal abundance, estimated with digital droplet PCR, was also higher in disease spots than visually healthy tissues (Table 2B and Figure S2). On the third date (July), the pathogen represented 65.5% of all fungal sequences. It was slightly more abundant in disease spots and in visually healthy tissues (Figure S1A), but this difference was not significant. Moreover, the total fungal abundance was similar in disease spots and visually healthy tissues (Figure S2), suggesting that the pathogen was widely dispersed on the third date.

**Table 1.** **Most abundant fungal species in grapevine foliar samples visually infected or not by the pathogen *Erysiphe necator*.** The relative abundances (RA, in % of sequences) and ranks of species were calculated for all leaf samples (TOTAL; n = 276) and for samples collected from visually healthy leaf blade of non-infected leaves (HNI; *n =* 93), visually healthy leaf blade of infected leaves (HI; *n =* 92) leaves, and disease spots (DI; *n* = 91) (Figure 1). Fungal sequences not assigned at the species level were not taken into account.

| Species | TOTAL |  | HNI |  | HI |  | DI |  |
| --- | --- | --- | --- | --- | --- | --- | --- | --- |
|  | Rank | RA | Rank | RA | Rank | RA | Rank | RA |
| *Erysiphe necator* | 1 | 35.1 | 3 | 14.9 | 1 | 22 | 1 | 68.9 |
| *Mycosphaerella tassiana* | 2 | 17.8 | 1 | 26.2 | 2 | 20.2 | 3 | 7 |
| *Filobasidium wieringae* | 3 | 13.9 | 2 | 15.5 | 3 | 19.8 | 4 | 6.2 |
| *Sporobolomyces roseus* | 4 | 9.2 | 4 | 9.9 | 4 | 10.3 | 2 | 7.4 |
| *Udeniomyces pyricola* | 5 | 4.4 | 5 | 6.3 | 5 | 5.3 | 5 | 1.6 |
| *Vishniacozyma victoriae* | 6 | 1.9 | 7 | 1.9 | 6 | 2.6 | 6 | 1.2 |
| *Erysiphe euonymicola* | 7 | 1.6 | 6 | 2.3 | 7 | 1.7 | 8 | 0.6 |
| *Dioszegia hungarica* | 8 | 1.2 | 8 | 1.6 | 8 | 1.3 | 9 | 0.6 |
| *Itersonilia pannonica* | 9 | 1 | 10 | 1.2 | 9 | 1.2 | 7 | 0.7 |
| *Symmetrospora coprosmae* | 10 | 0.8 | 11 | 1.2 | 11 | 1 | 16 | 0.2 |

**Table 2.** **Effects of date, cropping system (CS) and tissue type (TT) on (A) *E. necator* relative abundance, (B) total fungal abundance and (C) fungal community composition of grapevine leaves.** Foliar samples were collected on three dates (June 1, 23 and July 5), in two cropping systems (with cover crop or without cover crop), and in three tissue types (visually healthy leaf blade of non-infected leaves, visually healthy leaf blade of infected leaves, and disease spots; Figure 1). Linear models were used to analyze variations in pathogen abundance and total fungal abundance, while permutational analyses of variance were used to analyze community composition. Significant effects (p < 0.05) are in bold.

| Variable | Df | A. *E. necator*<br>relative abundance |  |  | B. Total fungal<br>abundance |  |  | C. Fungal community<br>composition |  |  |
| --- | --- | --- | --- | --- | --- | --- | --- | --- | --- | --- |
|  |  | F | Pr(>F) |  | F | Pr(>F) |  | F | R2 | Pr(>F) |
| Date | **2** | **55.29** | **1.00E-20** |  | **54.27** | **2.10E-20** |  | **5.1** | **0.0336** | **0.001** |
| CS | 1 | 0.1 | 7.50E-01 |  | 0.76 | 3.90E-01 |  | **1.5** | **0.005** | **0.021** |
| TT | **2** | **70.82** | **3.10E-25** |  | **50.87** | **2.40E-19** |  | 1 | 0.0068 | 0.377 |
| Date x CS | 2 | 0.29 | 7.50E-01 |  | 1.42 | 2.40E-01 |  | **9.7** | **0.0642** | **0.001** |
| Date x TT | **4** | **6.89** | **2.80E-05** |  | **17.9** | **5.50E-13** |  | 1.1 | 0.0148 | 0.13 |
| CS x TT | 2 | 1.08 | 3.40E-01 |  | 0.34 | 7.10E-01 |  | 1.2 | 0.0079 | 0.114 |
| Date x CS x TT | 4 | 1.11 | 3.50E-01 |  | 1.72 | 1.50E-01 |  | 1.1 | 0.0148 | 0.13 |
| Residuals | 258 |  |  |  | NA | NA |  | NA | 0.8529 | NA |
| **Total** | 275 |  |  |  |  |  |  | NA | 1 | NA |

### Infection does not destabilize the fungal community (no Anna Karenina effect)

In contrast to the expectations of hypothesis H1, fungal community composition was not found to differ between visually healthy tissues and disease spots, irrespective of the date. The composition changed significantly between cropping systems and, to a larger extent, between sampling dates (Table 2C and Figure S3). Community heterogeneity differed significantly among tissue types (ANOVA: df = 8; F = 37.7; p < 0.01), but contrary to the expectations of the Anna Karenina principle, community heterogeneity was not higher between infected tissues than between visually healthy ones. On the first sampling date, community heterogeneity was significantly lower in disease spots (Figure S1B), although the abundance of the pathogen was markedly higher (Figure S1A). The same results were obtained when *E. necator* was removed from the fungal community data (not shown), confirming the absence of Anna Karenina effect.

### Inferred microbial networks give relevant insights into the pathobiome

In accordance with hypothesis H2, we were able to show experimentally that some fungal associations inferred from metabarcoding data and environmental covariates corresponded to ecological interactions. The fungal association network inferred using the PLN model (Chiquet et al. 2018; 2019) was composed of 702 statistical associations between 61 fungal species, selected based on their prevalence (Figure S4A). Within this network, *E. necator* was negatively associated with 15 species and positively associated with 8 species (Figure 2); these 23 species were considered as the putative pathobiome of *E. necator*. All associations between *E. necator* and the pathobiome species, except two, were robust to subsampling (stability over 0.5; Table S1). None of these associations were known as ecological interactions in the literature. Literature-mining identified 9 abstracts in which *E. necator* and at least one species of its putative pathobiome were mentioned (Table S2), but these articles provided no experimental evidence of an interaction between the species. As a partial validation of the putative pathobiome of *E. necator* (Figure 2), we analyzed experimentally its interactions with 3 yeast species, *Buckleyzyma aurantiaca*, *Filobasidium wieringae* and *Vishniacozyma victoriae*, which were negatively associated with the pathogen in the PLN network (Figure 2). Microbiological experiments supported the antagonistic effect of *B. aurantiaca* on *E. necator*. The centrifuge supernatant from a *B. aurantiaca* liquid culture reduced the growth of *E. necator* by about 30% (Table 3) and significantly increased the number of collapsed conidia (Table S3 and Figure S5), when applied as a curative treatment. No effect was observed for the two other yeast species.

**Figure 2.**
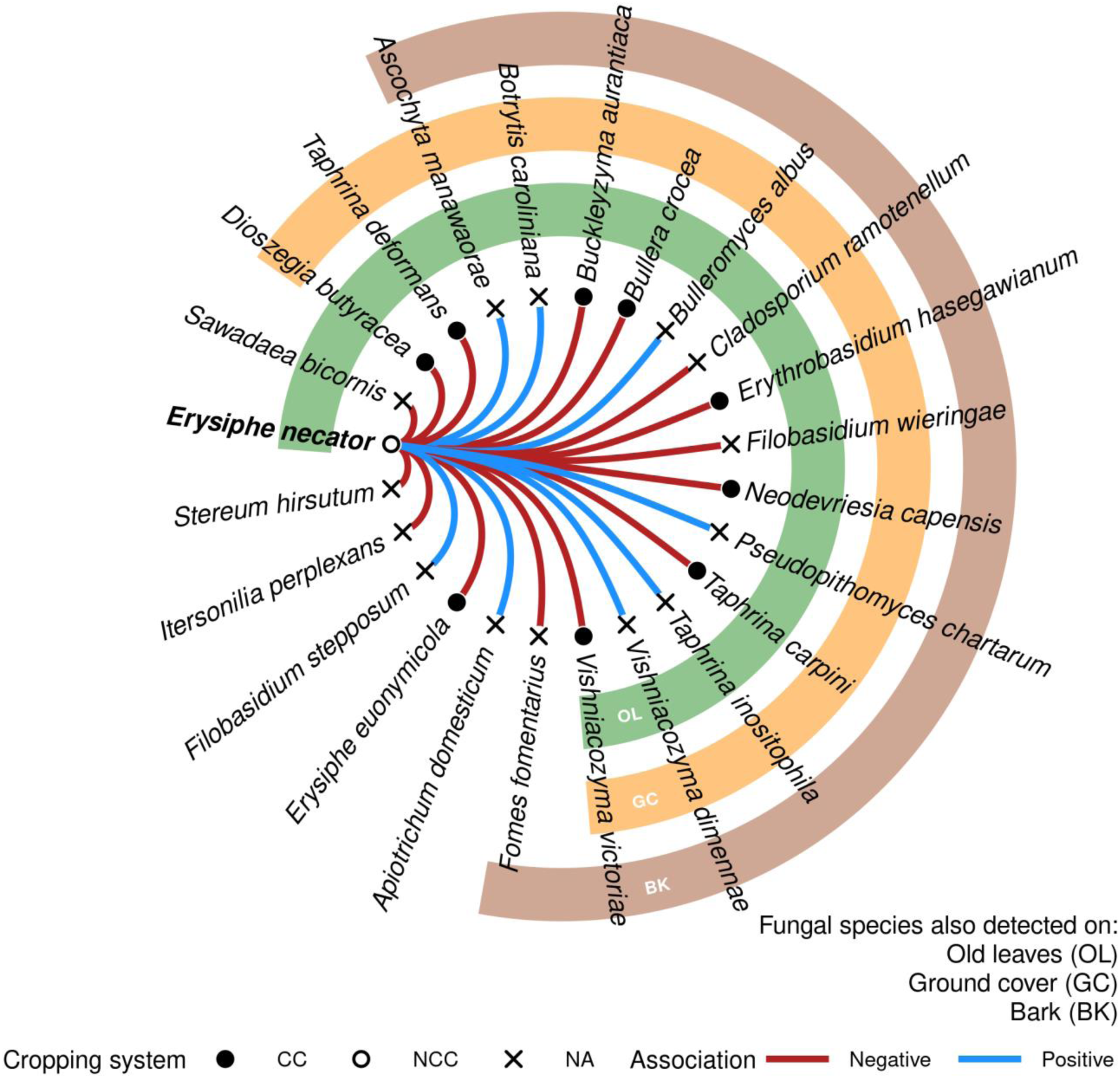
**Putative pathobiome of grapevine powdery mildew, *Erysiphe necator***, consisting of 25 species with positive (blue link) or negative (red link) associations with the pathogen. Node shape indicates whether these species were significantly favored by cover cropping (black circles), by the absence of cover (white circle) or neither (cross). Green, yellow and brown circles indicate whether the species were detected in other microbial environments (old leaves, ground cover and bark, respectively). The interactions between *E. necator* and 3 potential antagonists (*B. aurantiaca*, *F. wieringae* and *V. victoriae*) were tested experimentally.

**Table 3.** **Percent increase in *Erysiphe necator* (*En*) growth triggered by preventive or curative treatments with yeast strains in controlled conditions.** Significant effects of the treatments (p < 0.05) are in bold. Benzothiadiazole (BTH) is a growth inhibitor that was used as a positive control.

|  | Preventive treatment |  |  |  | Curative treatment |  |  |  |
| --- | --- | --- | --- | --- | --- | --- | --- | --- |
|  | Pellet |  | Supernatant |  | Pellet |  | Supernatant |  |
| Yeast strain | En growth (in %) | p-value | En growth (in %) | p-value | En growth (in %) | p-value | En growth (in %) | p-value |
| Control (BTH) | **-98.62** | **0** | **-90.62** | **0** | **-85.62** | **1.70E-14** | **-86.25** | **0** |
| *B. aurantiaca* | -1.87 | 0.33 | 2.25 | 0.99 | -19.63 | 0.18 | **-31.25** | **0** |
| *U. pyricola* | -1 | 0.85 | 6 | 0.83 | **-32.5** | **0** | -3.75 | 0.97 |
| *V. victoriae* | -0.37 | 0.99 | 7.63 | 0.67 | -10 | 0.79 | 3.88 | 0.97 |
| *D. hungarica* | 0.25 | 1 | 0.63 | 1 | -5 | 0.98 | -10.62 | 0.44 |
| *F. wieringae* | 0.5 | 0.98 | 6.38 | 0.8 | -10.63 | 0.75 | -1.87 | 1 |
| *S. roseus* | 0.88 | 0.9 | 0.13 | 1 | -6.88 | 0.93 | -5.25 | 0.91 |
| *F. oeirense* | 1.13 | 0.78 | -4.5 | 0.93 | -19.38 | 0.19 | -0.87 | 1 |
| *C. macerans* | 1.38 | 0.64 | 6.25 | 0.81 | -10.63 | 0.75 | 5 | 0.93 |

We also analyzed experimentally the interactions between *E. necator* and 5 additional yeast species (*Cystofilobasidium macerans*, *Dioszegia hungarica*, *Filobasidium oeirense*, *Udeniomyces pyricola* and *Sporobolomyces roseus*) and all interactions between the 8 yeast species. All the yeast species were among the most abundant species in grapevine leaves and five belonged to the top ten (Table 1). We found several discrepancies between the PLN association network and the ecological interactions revealed by the co-cultures. Spot-on-lawn experiments (Figure S4C) revealed 3 growth-promoting interactions and 12 growth-inhibiting interactions among the yeast species (Figure S4D). This *in vitro* interaction network shared only two links with the PLN association network (Figure S4B), between *C. macerans* and *U. pyricola* and between *F. oeirense* and *C. macerans*. The positive association between *C. macerans* and *U. pyricola* was upheld by the spot-on-lawn assay as *U. pyricola* enhanced the growth of *C. macerans*. In contrast, the link between *F. oeirense* and *C. macerans* was found as a positive association with PLNnetwork, while the growth of *C. macerans* was inhibited by *F. oeirense* in the spot-on-lawn assay. Similarly, although there was no association between *U. pyricola* and *E. necator* in the PLN network, we found that the pellet of *U. pyricola* culture inhibited *E. necator* growth when applied as a curative treatment (Table 3).

### Cropping system influences the abundance of pathobiome members

Cover cropping increased significantly the abundance of 9 species belonging to the putative pathobiome of *E. necator*, including the yeast *B. aurantiaca* (Figure 2 and Table S4). These 9 species were putative antagonists of *E. necator* and eight of them, including *B. aurantiaca*, were generalist fungal species found in grapevine leaves but also in bark and ground cover (Figure 2 and Table S1). These findings suggest that cropping systems modulate pathobiomes, in accordance with hypothesis H3. Interestingly, only the pathogen *E. necator* was significantly favored by the absence of a cover crop according to differential abundance analyses (Table S4). This pattern, which might be accounted for by the significant increase in vine vigor in the absence of cover crop (Figure S6A), was however not confirmed by the visual disease severity assessment performed mid-July (Figure S6B).

## DISCUSSION

Plant pathologists have long believed that diseases result from interactions between pathogens, their hosts and the abiotic environment (the so-called ‘disease triangle’). A paradigm shift occurred in 2014 with the integration of the microbiota into this framework (Vayssier-Taussat et al. 2014; Kemen et al. 2015; Vandenkoornhuyse et al. 2015). The new paradigm led to the concept of the pathobiome, which corresponds to all the microorganisms associated with a host that regulate disease (Vayssier-Taussat et al. 2014; Brader et al. 2017; Bass et al. 2019). Contemporaneous meta-barcoding and meta-genomic approaches revealed the huge microbial diversity associated with soil and plants, including grapevine (Zarraonaindia et al. 2015). This paradigm shift increased the complexity of the studied systems and triggered an increase in the use of network approaches in plant pathology. To identify the microorganisms protecting plants against pathogens, several authors suggested considering the statistical associations within meta-barcoding datasets as a proxy for microbial interactions (Poudel et al. 2016; Vacher et al. 2016; Garett et al. 2018; Vannier et al. 2019), with the ultimate goal of identifying (groups of) microbial taxa that compete for space or resources with pathogens, parasitizes pathogens or have an antibiosis activity against them (Massart et al. 2005, Abdelfattah et al. 2018). In this study, we assessed the relevance of this novel line of research using grapevine powdery mildew (*Erysiphe necator*; Gadoury et al. 2012, Armijo et al. 2016) as a model system.

We analyzed the microbiota of both healthy and diseased foliar tissues to discover microbial species with potential antagonistic activity against grapevine powdery mildew. We then used a new statistical approach, the PLNnetwork variant of the PLN model family (Chiquet et al. 2018; 2019), to generate novel hypotheses of microbial interactions in the form of a network of positive and negative associations between microbial species. The network was inferred from a cross-sectional meta-barcoding dataset including a large number of samples (276 in total) and several environmental variables associated with each sample. The number of samples was in line with the recent recommendation by Hirano and Takemoto (2019) of at least 200 samples per network, and the inferred associations were robust. Network analysis suggested that the pathogen species was associated with 23 other fungal species, forming its putative pathobiome. We tested some of the hypotheses of interaction using co-cultures. Interestingly, we found that the yeast *Buckleyzyma aurantiaca* reduced pathogen growth and altered the shape of asexual reproduction organs, the conidia. These findings upheld the negative association between the abundances of *B. aurantiaca* and *E. necator* revealed by the network approach, and make this yeast a potential biocontrol candidate deserving further investigation. *B. aurantiaca* (ex *Rhodotorula aurantiaca*) has previously been reported as a leaf-colonizing yeast which, when isolated from disease-free plants, was able to protect melon plants against bacterial fruit blotch (de Melo et al. 2015). A hypothesised association involving two yeast species, *Cystofilobasidium macerans* and *Udeniomyces pyricola*, was also supported by co-cultures. These findings corroborate our expectation that network inference methods can detect signals of ecological interactions between microorganisms (Das et al. 2018; Durán et al. 2018; Gao et al. 2018) and are useful tools for developing future microbiome-based crop protection strategies.

Our results also suggested that cover cropping promotes disease control. We found that cover cropping increased the abundance of *B. aurantiaca* and eight other putative antagonists of *E. necator*. All but one of these species were detected in the ground cover and on vine leaves. Cover cropping decreased significantly both the vigor of the vines and the abundance of the pathogen in the molecular dataset, confirming previous results (Valdés-Gómez et al. 2011). Hence, our findings suggest that cover cropping might regulate powdery mildew by both influencing vine physiology (Valdés-Gómez et al. 2011) and increasing the abundance of generalist fungal species that compete with *E. necator*. The same mechanisms could account for the reduced severity of black rot and downy mildew (Figure S6) in the presence of cover crops. Cover cropping is already widely used in vineyards to protect the soil, limit herbicide use (Pertot et al. 2017) and strengthen natural pest regulation by favoring predatory insects (Saenz-Romo et al. 2019). Our study provides additional support for the potential benefits of ground cover via the maintenance of generalist fungal species that can compete with foliar pathogens.

Finally, our analyses revealed what distinguishes the microbiota of visually healthy tissues from that of infected tissues, which is an important step for understanding disease ecology and developing microbiome-based management strategies. Surprisingly, *E. necator* was among the dominant species of grapevine leaves in our study, even in visually healthy tissues. The relative abundance of the pathogen was much higher in our study (between 15 to 70% depending on the tissue type) than in other studies of grapevine foliar fungal communities, where it represented less than 5% (Fort et al. 2016; Gobbi et al. 2020). Several factors, including leaf age and sampling date, might account for this difference as there is marked turnover in the grapevine foliar fungal community along the vegetative season (Pinto & Gomes 2016; Fort et al. 2016; Gobbi et al. 2020). The artificial inoculation of the pathogen on some vines at the beginning of the vegetative season is also probably responsible for the higher relative abundance of the pathogen in our study. As expected, the pathogen was significantly more abundant in disease spots than in visually healthy tissues of the same leaf, and was more abundant in visually healthy tissues of leaves harboring disease spots than in leaves that appeared completely healthy, except for the final sampling date when the abundance of the pathogen was high in all tissue types. The presence of pathogen spores on visually healthy tissues could account for this latter result. Moreover, we found that the composition of the fungal community was similar in visually healthy tissues and disease spots, in contrast to the results previously obtained for oak powdery mildew, *Erysiphe alphitoides* (Jakuschkin et al. 2016). This result would suggest that *E. necator* interacts with only a few fungal species, as revealed by the network approach, but that these interactions do not shape the whole fungal community. Furthermore, we found no evidence to support the Anna Karenina principle, which predicts higher heterogeneity in microbial community composition in infected tissues (Zaneveld et al. 2017). This suggests that the Anna Karenina principle is not ubiquitous and that the theoretical framework guiding pathobiome research needs further resolution. The absolute fungal load, which was estimated using microfluidic quantitative PCR, varied between visually healthy and infected tissues, in contrast to the composition of the fungal community. On the second sampling date, fungal load was significantly higher in infected tissues than in visually healthy tissues, corroborating the expectation that microbial absolute load is an important indicator of dysbiosis (Karasov et al. 2019). We suggest that the absolute quantification of microbial taxa abundance, which is often neglected in microbial community ecology studies based on meta-barcoding approaches, is necessary for understanding the pathobiome. Absolute quantification can be performed by coupling metabarcoding community data with total abundance data (Dannemiller et al. 2014; Vandeputte et al. 2017; Props et al. 2017), developing microfluidic quantitative chips targeting pathobiome members (Kleyer et al., 2017; Crane et al., 2018) and by using meta-genomic approaches (Karasov et al. 2019). These approaches will improve the reliability of pathobiome networks because they help reduce compositionality bias, i.e. false associations generated by the comparison of relative abundances (Röttjers & Faust 2018).

Overall, this study demonstrates that microbial networks give valuable insights into plant pathobiomes, can guide the search for novel biocontrol agents and suggest disease management approaches, such as the cover cropping systems shown here. To complete the pathobiome network of grapevine powdery mildew, future studies should collect data on the bacterial, oomycete, virus and even insect components of the vine community because inter-kingdom interactions may influence the structure and dynamics of microbial networks (Agler et al. 2016; Jakuschkin et al. 2016; Durán et al., 2018; Tipton et al. 2018). This additional information will also improve the inference of associations because missing species can generate apparent and indirect associations (*e.g.* Li et al. 2016). The integration of prior information for certain interactions (reviewed by Panstruga & Kuhn 2019 for powdery mildews) will also improve network inference (Lo and Marculescu 2017). The analysis of microbial networks associated with wild relatives of cultivated grapevine or hybrids could also be relevant, given that a recent study highlighted higher abundance of beneficial symbionts in wild vines (Kernaghan et al. 2017). Advances in culturomics (Lagier et al. 2018) and synthetic microbial communities (Kehe et al. 2019; Vannier et al. 2019) will also undoubtedly benefit the field of pathobiome research. In the present study, we did a partial validation of the pathobiome network using specific microbial cultures. In the future, microbiological experiments will enable the comparison and calibration of network inference methods based on synthetic microbial networks, which will allow us to select the method and the parameters of network reconstruction best suited to the features of the microbial system.

## EXPERIMENTAL PROCEDURE

### Study site and sampling design

The sampling campaign occurred in spring and summer 2016 in an experimental vineyard located near Bordeaux (INRA, Villenave d’Ornon, France; 44°47’24.0”N 0°34’33.6”W; Figure 1A). This experimental vineyard was planted in 1991 with *Vitis vinifera* L. cv. Merlot grafted onto 101-14 rootstock at a density of 5350 vines ha^-1^ (1.7m x 1.1m). Sampling was performed in a sub-plot of 5 largely untreated rows (Figure 1B) where very few chemical pesticides had been applied over the last ten years, and none in the last four years (Table S5). This sub-plot was composed of two cropping systems (Figure 1C): (i) perennial cover crop in the inter-rows (i.e. cover crop: CC) and (ii) chemical weed control with glyphosate (i.e. no cover crop: NCC). We defined eight experimental units across the sub-plot: four units in NCC areas and four units in CC areas, an experimental unit being defined as a group of 5 adjacent vines in the same row (Figure 1D).

On April 22, we inoculated the central vine of each experimental unit with *E. necator* (Figure 1E). The inoculum was a monoconidial isolate (strain S16) collected that year in a greenhouse (INRA Villenave d’Ornon, Bordeaux, France) and bulked on Cabernet Sauvignon leaves from greenhouse grown cuttings (Cartolaro and Steva 1990). Leaves were then regularly checked for powdery mildew colonies. The first symptoms appeared on May 5 for the inoculated leaves, and May 18 for other leaves. Leaf age was monitored in all experimental units based on biweekly records of newly grown leaves. In addition, a weather station located on the edge of the vineyard allowed us to monitor local variations in air temperature and humidity throughout the experiment.

Grapevine leaves were sampled on June 1, June 23 and July 5 (40, 62 and 77 days post-inoculation (dpi), respectively) (Figure 1F). On each date, we collected four infected (I) and four non-infected (NI) leaves of approximately 20-day old in each experimental unit by visual assessment (Figure 1G). In total, we collected 192 leaves, corresponding to 8 leaves × 3 dates × 8 experimental units. Based on current knowledge of the powdery mildew cycle (Calonnec et al. 2006, 2009), all sampled leaves had received secondary inoculum at their optimum leaf age susceptibility, i.e. between 6 to 9 days old (Calonnec et al. 2018). They were also old enough to harbor disease symptoms, knowing that the average latency period is between 7 and 10 days and that the symptoms are apparent 3-4 days after the beginning of sporulation. The percentage of lower leaf surface covered with powdery mildew was visually assessed for each visually infected leaf. We also measured the distance between each sampled leaf and three potential environmental sources of microorganisms: the ground, the cordons and the artificially inoculated leaf in the same experimental unit.

We placed leaves in individual sterile plastic bags (Whirl-Pak®, USA) and took them to the laboratory in a cooler with ice. We processed the leaves on the day of sampling in the sterile environment of a MICROBIO electric burner (MSEI, France). Tissues were collected from visually healthy leaf blade of non-infected leaves (HNI), visually healthy leaf blade of infected leaves (HI) and disease spots (DI) (Figure 1E). We collected two foliar discs of 6 mm diameter in each tissue type available and then placed the two disks together in a collection microtube of a 96-well plate (QIAGEN), with two autoclaved glass beads. The processed leaves were stored on ice in individual closed plastic bags to avoid water loss until we had measured leaf fresh weight and surface area using the WinFOLIA® software (Regent Instrument, Canada). Dry weight was measured after drying the leaves in an oven at 65°C for 72h.

On July 5, additional environmental samples were collected in each experimental unit. We collected two old leaves (approximately 70 days old) from the central vine of each experimental unit, close to the place where the inoculation had been performed, and placed them in a sterile plastic bag (Whirl-Pak®, USA). The inoculated leaf was also collected if it was still attached to the vine. We collected small pieces of bark (including mosses and lichens) on three vines of each experimental unit and stored them in 50mL sterile Falcon tubes. Finally, we collected fragments of ground cover (including the upper soil) beneath three vines of each experimental unit, and these were stored in 50mL sterile Falcon tubes.

On July 11, we assessed the sub-plot phytosanitary status and canopy vigor. The severity (% of diseased lower leaf surface) of four diseases, *i.e.* powdery mildew (*E. necator*), downy mildew (*Plasmopara viticola*), black rot (*Guignardia bidwellii*) and grape erineum mite (*Colomerus vitis*), was visually evaluated on 12 randomly chosen leaves on each vine of each experimental unit. We estimated vine canopy vigor as the product of the number of shoots per vine by the number of leaves on the longest shoot.

### DNA extraction and sequencing

Leaf discs were cold ground at 1500 rpm with the Geno/Grinder® twice for 30 s, with manual shaking between each grinding step. Then, plates were centrifuged for 1 min at 6200 rpm. Total DNA was extracted using the DNeasy Plant Mini Kit (Qiagen, USA) following the manufacturer’s instructions. The ITS1 region of the fungal ITS rDNA gene (Schoch et al. 2012) was amplified using the primers ITS1F-ITS2 (White et al. 1990; Gardes and Bruns 1993). To avoid a two-stage PCR protocol, each primer contained the Illumina adaptor sequence and a tag (ITS1F: 5’-CAAGCAGAAGACGGCATACGAGATGTGACTGGAGTTCAGACGTGTGCTCTTCCGATCTxxxxxx xxxxxxCTTGGTCATTTAGAGGAAGTAA-3’; ITS2: 5’-AATGATACGGCGACCACCGAGATCTACACTCTTTCCCTACACGACGCTCTTCCGATCTxxxxxxx xxxxxGCTGCGTTCTTCATCGATGC-3’, where “x” is the 12 nucleotides tag). The PCR mixture (20 µL of final volume) consisted of 10 µL of 2X QIAGEN Multiplex PCR Master Mix (1X final), 2 µL each of the forward and reverse primers (1 µM), 4 µl of water, 1 µL of 10 mg.ml^-1^ BSA and 1 µL of DNA template. PCR cycling reactions were conducted on a Veriti 96-well Thermal Cycler (Applied Biosystems) using the following conditions: initial denaturation at 95°C for 15 min followed by 35 cycles at 94°C for 30 s, 57°C for 90 s, 72°C for 90 s with final extension of 72°C for 10 min. ITS1 amplification was confirmed by electrophoresis on a 2% agarose gel. Each PCR plate contained one negative extraction control, one negative PCR control, and one positive control. The negative extraction control corresponded to an empty microtube left on the collection plate. The negative PCR control corresponded to PCR mix without any DNA template. The positive PCR control was composed of an equimolar mixture of the DNA of two marine fungal strains, *Candida oceani* and *Yamadazyma barbieri* (Burgaud et al. 2011, 2016). These strains were chosen as positive controls as they were unlikely to be found in or on grapevine leaves. PCR products were purified, quantified (Quant-it PicoGreen dsDNA assay kit; Thermo Fisher Scientific) and equimolarly pooled (Hamilton Microlab STAR robot). Average size fragment was checked using Tapestation instrument (Agilent Technologies). Libraries were sequenced on the MiSeq Instrument (Illumina) with the reagent kit v2 (500-cycles). Sequence demultiplexing (with exact index search) was performed at the PGTB sequencing facility (Genome Transcriptome Facility of Bordeaux, Pierroton, France) using DoubleTagDemultiplexer. Additional environmental samples collected on July 5 were processed separately following the protocols described in Fort et al. (2019). They were manually ground in liquid nitrogen, amplified using a two-step PCR and then sequenced on a separate MiSeq run at the GetPlaGe sequencing facility (Toulouse, France).

### Fungal DNA quantification

Fungal DNA was quantified with digital droplet PCR assays (ddPCR™; Hindson et al. 2011) using the universal fungal primer pair ITS1F (5’-TCCGTAGGTGAACCTGCGG-3’) and 5.8S (5’-CGCTGCGTTCTTCATCG-3’) validated by Fierer et al. (2005). Assays were carried out with the QX200 Droplet Digital PCR (ddPCR™) System from Bio-Rad at the PGTB sequencing facility (Pierroton, France). The PCR reactions were carried out in a final volume of 20 μl using the ddPCR™ EvaGreen Supermix (Bio-Rad, USA). The reaction mix consisted of 10 µl of 2X EvaGreen Supermix, 2.5 μL of each primer at 1.2 μM and 3 μL of DNA template or ultrapure water in the negative control. The mix containing the sample was partitioned into droplets with the QX200 Droplet Generator and then transferred to 96-well PCR plates. A thermocycling protocol [95 °C × 5 min; 40 cycles of (95°C × 30 s, 53°C × 1 min 30 s), 4 °C × 5 min, 90°C × 5 min] was undertaken in a Bio-Rad C1000 (Bio-Rad, USA). QX200 droplet reader analyzed each droplet individually to detect the fluorescence signal. The number of copies of the target DNA sequence per μl of sample was determined from the number of positive droplets (out of an average of ∼20k droplets per sample) estimated from fluorescence signals of both samples and negative controls using the Umbrella procedure implemented in R (Jacobs et al. 2017). Total fungal abundance was then obtained by multiplying the obtained concentration by the mix volume and after adjusting for the 1/100 dilution of the DNA extract.

### Bioinformatic analysis

We used DADA2 v1.8 (Callahan et al. 2016) to describe fungal communities in terms of Amplicon Sequence Variants (ASVs, Callahan et al. 2017) because bioinformatic approaches based on DADA2 have been found to recover accurately the composition of an artificial fungal community (Pauvert et al. 2019). We followed the DADA2 ITS workflow (https://benjjneb.github.io/dada2/ITS_workflow.html) except on the read assembly step. Only forward reads in which the primer sequence was found exactly by cutadapt (Martin 2011) were processed with DADA2. Quality filtering retained sequences with less than one expected error and longer than 50 bp. ASVs were subsequently inferred for each sample. Chimeric sequences were then identified using the consensus method of the *removeBimeras* function. ASVs identical in sequence but not in length were finally combined with the *collapseNoMismatch* function. Taxonomic assignments were then performed using the RDP classifier (Wang et al. 2007) implemented in DADA2 and trained with the latest UNITE database (UNITE Community 2019). ASV table and taxonomic assignments were imported in R through the phyloseq package (McMurdie et al. 2013). Only ASVs assigned to a fungal phylum were kept.

Positive and negative controls were used to remove contaminants, as described by Galan et al. 2016. The ASVs corresponding to positive control strains (*Candida oceani* and *Yamadazyma barbieri*) were identified by aligning the ASV sequences with the Genbank reference sequences of control strains (*C. oceani* KY102240 and *Y. barbieri* LT547714) using a similarity threshold of 100% with *usearch_global* function of VSEARCH (Rognes et al. 2016). The cross-contamination threshold (T_CC_) was defined as the maximal number of sequences of each ASV found in negative or positive control samples. The false-assignment threshold (T_FA_) was defined as T_FA_ = N × R_fa_ where R_fa_ is the highest sequence count of a positive control strain in a non-control sample, divided by the total number of sequences of the strain in the whole run and N is the total number of sequences of each ASV. ASVs were removed from all samples where they harbored fewer sequences than either threshold (T_FA_ or T_CC_) using a custom script (https://gist.github.com/cpauvert/1ba6a97b01ea6cde4398a8d531fa62f9). The ASV table was finally aggregated at the species level and ASVs that could not be assigned at the species level were removed. All analyses were performed on the resulting species × sample matrix.

### Hypothesis testing

All statistical analyses were performed in R (R core Team 2018). To validate the sampling design, we first checked that disease spots (DI) had a higher fungal load and a higher relative abundance of *E. necator* than visually healthy tissues (HNI and HI). Variations in fungal total abundance (log-transformed) and in *E. necator* sequence counts (CLR-transformed) were analyzed using linear regressions performed with the *lm* function. The models had tissue type (HNI, HI or DI), cropping system (CC or NCC), sampling date (40, 62 or 77 dpi) and their interactions as fixed effects. *F*-tests were used to assess the significance of the fixed effects and post-hoc pairwise comparisons were performed for the significant interactions using Tukey’s adjustment method with the emmeans R package (Lenth 2018). Then, we tested hypotheses H1, H2 and H3.

#### H1: Successful infection events destabilize plant-associated microbial communities and increase their heterogeneity (Anna Karenina principle)

To test hypothesis H1, we investigated whether infection altered the composition of foliar fungal communities using permutational analysis of variance (PERMANOVA; Anderson et al. 2001) performed with the *adonis* function of the vegan R package (Oksanen et al. 2018). The model had tissue type (HNI, HI or DI), cropping system (CC or NCC), sampling date (40, 62 or 77 dpi) and their interactions as fixed effects. The analysis was performed using CLR-transformed community data, as advised by Gloor et al. (2017). A total of 128 CLR-transformed species × sample matrices were generated using the aldex.clr function of ALDEx2 (Fernandes et al. 2014) and averaged. Euclidean distances among samples of the average matrix were then used to test differences in community composition among conditions. These analyses were performed twice, with and without *E. necator*.

We then investigated whether compositional heterogeneity was larger among diseased samples (DI) than among healthy ones (HI and HNI), by performing tests of homogeneity of multivariate dispersion (Anderson 2006) with the *betadisper* function of the R vegan package (Oksanen et al. 2018). We defined community heterogeneity within a group of samples as the average distance between samples and the group centroid, and calculated it for every combination of sampling date and tissue type. Differences in heterogeneity between groups were tested using *F*-tests. Post-hoc pairwise comparisons were performed with Tukey’s adjustment method with the emmeans R package (Lenth 2018).

#### H2: Interactions among microorganisms within pathobiomes can be detected by inferring microbial networks from metabarcoding data and environmental covariates

To test hypothesis H2, we inferred the foliar microbial network the species × sample matrix and then tested some of the associations using co-culture experiments and text-mining.

##### Network inference

Species included in the network reconstruction were selected based on their prevalence (Röttjers & Faust, 2018; Cougoul et al. 2019a). Only species present in more than 20% of the samples of any experimental unit were kept. Joint variations in sequence counts between fungal species were modelled with the PLNmodels R package (Chiquet et al, 2019) to account for uneven sequencing depth among samples and for environmental covariates potentially explaining covariation in fungal species abundances across samples. Direct statistical associations between species were inferred with the PLNnetwork function. First, the extended BIC criteria (Chen and Chen 2008) was used to perform model selection among a 40-size grid of penalties controlling the sparsity of the underlying network (Chiquet et al. 2018; 2019). Then, the stability of each association of the selected network was calculated as its selection frequency in the bootstrap subsamples of the StARS procedure (Liu et al. 2010). Environmental covariates included several foliar traits and climate variables that may influence fungal development: leaf age, leaf water content, specific leaf area and average vapor pressure deficit experienced by the leaf since unfolding. These covariates were introduced in the model to get rid of fungal species associations triggered by similar habitat requirements. The distance between the sampled leaf and environmental sources of microorganisms was also included as a covariate to get rid of fungal associations triggered by joint colonization events. Only the distance between the sampled leaf and old leaves was introduced in the model as the three distances measured were significantly correlated. To account for additional, non-measured environmental variations associated with the experimental design, we also included the three main factors of the experiment (sampling date, experimental unit and tissue type) as covariates. Finally, sequencing depth (log-transformed) was included as an offset in the model to control for spurious associations triggered by compositionality.

##### Network validation using co-cultures

We then used co-cultures to test whether the statistical associations among fungal species revealed by the PLN model corresponded to ecological interactions. We tested the pairwise interactions among eight yeast strains present in the PLN network. These eight yeast strains, which all belonged to the most abundant fungal species of the dataset, were bought from the CBS collection (Westerdjik Fungal Biodiversity Institute, Utrecht, The Netherlands). The strains were *Buckleyzyma aurantiaca* CBS 8074, *Cystofilobasidium macerans* CBS 9032, *Dioszegia hungarica* CBS 7091, *Filobasidium oeirense* CBS 8681, *Filobasidium wieringae* CBS 1937, *Udeniomyces pyricola* CBS 6754, *Sporobolomyces roseus* CBS 486 and *Vishniacozyma victoriae* CBS 9000. Strains were revived following the CBS instructions and further maintained in collection on MEA or PDA media [Malt Extract Agar (Malt 15 g.l^-1^, Agar 20 g.l^-1^); Potato Dextrose Agar, Biokar diagnostics, France (Potato extract 4 g.l^-1^, Glucose 20 g.l^-1^, Agar 15 g.l^-1^)]. To produce inoculum for the co-culture experiments, yeast strains were grown in liquid ME medium (Malt Extract, 15 g.l^-1^) at 22°C for 48h. The number of yeast cells was then counted using a hemacytometer before being homogenized at the same concentration. The eight yeast strains were confronted to one another using a spot-on-lawn assay (Polonelli et Morace, 1986; Addis et al., 2001). Inoculations were performed on MEA medium buffered at pH 4.5 with 0.5 M phosphate-citrate buffer (Heard and Fleet, 1987). Approximately 10^5^ cfu.ml^-1^ of each strain was suspended in 15 ml of sterile MEA medium (pH 4.5) maintained at 45°C and then poured into sterile Petri dishes. For each strain, five replicates were made, totaling 40 plates. Each of the eight strains was then drop-inoculated (50 µl at 10^5^ cfu.ml^-1^) onto the surface of a Petri dish. Once the drops dried, the plates were sealed with Parafilm® and incubated at 22°C. Yeast growth was visually assessed 4 and 7 days after inoculation. We considered that the MEA-included strain either inhibited or promoted the growth of a drop-inoculated strain, if the latter had either a reduced or increased growth relative to its own MEA-included confrontation.

Then we tested the pairwise interactions between the eight yeast strains and one isolate of the pathogen *E. necator*. This latter was a monoconidial isolate (strain S19) collected in 2019 in a greenhouse (INRA Villenave d’Ornon, Bordeaux, France) and bulked on Cabernet Sauvignon leaves (Cartolaro and Steva 1990). Liquid yeast cultures incubated in ME for 48h were centrifuged at 5000 rpm and 4°C for 15 min. Supernatant (liquid medium) and pellet (yeast cells) were then separated. Pellets were resuspended in sterile distilled water at a concentration of 10^6^ cfu.ml^-1^. Powdery mildew isolate S19 was inoculated under sterile conditions on sterilized grape leaf discs (eight 2-cm discs per plate, each coming from a different leaf). Both components, *i.e.* supernatants and resuspended pellets, were tested against *E. necator* in a preventive assay (applied 24h before *E. necator*) and a curative one (applied 6h after *E. necator*). In total, we inoculated 320 foliar discs, corresponding to 8 foliar discs per plate x 10 plates (8 plates, one for each yeast strain + 1 negative control + 1 positive control) x 2 assays (preventive and curative) x 2 components tested (medium supernatant and yeast cells). Negative controls were treated either with sterile distilled water or sterile ME respectively in the pellet and supernatant conditions. Positive controls were treated with BTH (S-methyl benzo[1,2,3]thiadiazole-7-carbothioate, Bion®, 50WG, Syngenta) at a concentration of 0.1% active ingredient, a growth inhibitor of powdery mildew (Dufour et al. 2013). After incubation for 12 days at 22°C with a 16 h day/8 h night photoperiod, *E. necator* growth was evaluated on each foliar disc by assessing visually the diseased leaf area (DLA, in %). The effect of each yeast strain on *E. necator* growth was estimated relative to the negative control using *trt.vs.ctrl* contrasts in emmeans (Lenth 2018). Moreover, the number of foliar discs with altered conidia in the presence of a yeast strain was measured and compared to that of the negative control using a Fisher exact test with the alternative = “greater” option.

##### Network validation using text-mining

Finally, we investigated whether the subset of the association network involving *E. necator* corresponded to interactions described in the literature. Using a custom R script (Methods S1), we investigated whether fungal species names associated in the network co-occurred in articles of the Scopus database (that includes article titles, abstracts, keywords and references) using its API through the rscopus R package (Muschelli 2019). Obligate and anamorph synonyms of fungal species names were queried from the MycoBank webservice (Robert et al. 2013). The resulting articles were searched for experimental evidence of ecological interaction between the selected species.

#### H3: Cropping systems influence the abundance of microorganisms forming pathobiomes

To test hypothesis H3, we investigated whether the fungal species statistically associated with *E. necator* in the PLN association network (i.e. its putative pathobiome) differed in abundance between CC and NCC cropping systems. ALDEx2 (Fernandes et al. 2014) was used to detect differentially abundant species as this method is suitable for compositional data (Gloor et al. 2017) and outputs very few false-positives with default values (Thorsen et al. 2016). Species with a false discovery rate below 0.1 after Benjamini-Hochberg adjustment were considered as differentially abundant. To unravel the ecology of the fungal species belonging to the pathobiome of *E. necator*, we searched for their presence in the environmental samples. Finally, to better understand the effects of cropping systems, we compared grapevine vigor and disease severity between CC and NCC experimental units using t-tests.

## ACKNOWLEDGEMENTS

We thank Jérôme Jolivet, Lionel Druelle, Hany El Manawy and Benoît Bénéteau for their help during the sampling campaigns, and Isabelle Demeaux for her help with *E. necator* experiments. We thank Gaétan Burgaud for kindly providing marine fungal strains used as positive controls. We also thank Gregory Gambetta, Guilherme Martins, Frédéric Barraquand, Isabelle Lesur, Adrien Rush and all members of the ANR NGB Consortium (ANR-17-CE32-0011) for useful comments on preliminary results. We thank the CLIMATIK portal (INRAE Avignon) for providing climatic data concerning the weather station 33550003. We thank the Genotoul sequencing facility (Get-PlaGe) and the PGTB sequencing facility for sequencing, and the Genotoul bioinformatics facility (Bioinfo Genotoul) for providing computing and storage resources. We thank the INRA MEM metaprogram (Meta-Omics of Microbial Ecosystems) for financial and scientific support (Learn-biocontrol project). Additional funding was received from the LABEX COTE (ANR-10-LABX-45), the LABEX CEBA (ANR-10-LABX-25-01), INRA EcoServ metaprogram on ecosystem services (IBISC project) and the Aquitaine Region (Athene project, n°2016-1R20301-00007218). The management of the experimental site was partly funded by the AFB (French Agency for Biodiversity) within the DEPHY network. CP’s PhD grant was funded by the INRA and Bordeaux Sciences Agro (BSA).

## AUTHORS CONTRIBUTION

CV, DAB and JV had the original idea for the project. JV and AC performed inoculations. JV, CV and AC performed leaf monitoring and sampling. JV and CV processed the sampled leaves. TF and JFA performed the extractions and amplifications. EC performed the sequencing. CP and MM quantified fungal abundance using ddPCR. JV and CP performed the microbiology experiments. CP performed the bioinformatic and statistical analyses. SR and JC provided methods and advice on network inference. CV and JV coordinated all stages of the work. CP wrote the first draft of the manuscript, and CV and JV made a major contribution to the writing of the final draft. All authors revised the manuscript.

## DATA AVAILABILITY

The raw sequence data for grapevine leaves and environmental microbial sources were deposited in Dataverse and are available at https://doi.org/10.15454/A24N4C and https://doi.org/10.15454/XXBU8Y, respectively. Bioinformatic and statistical analysis scripts are available at https://doi.org/10.15454/WLHBP6 and https://doi.org/10.15454/5WD6P6, respectively.

## SUPPLEMENTARY MATERIALS

**Figure S1.**
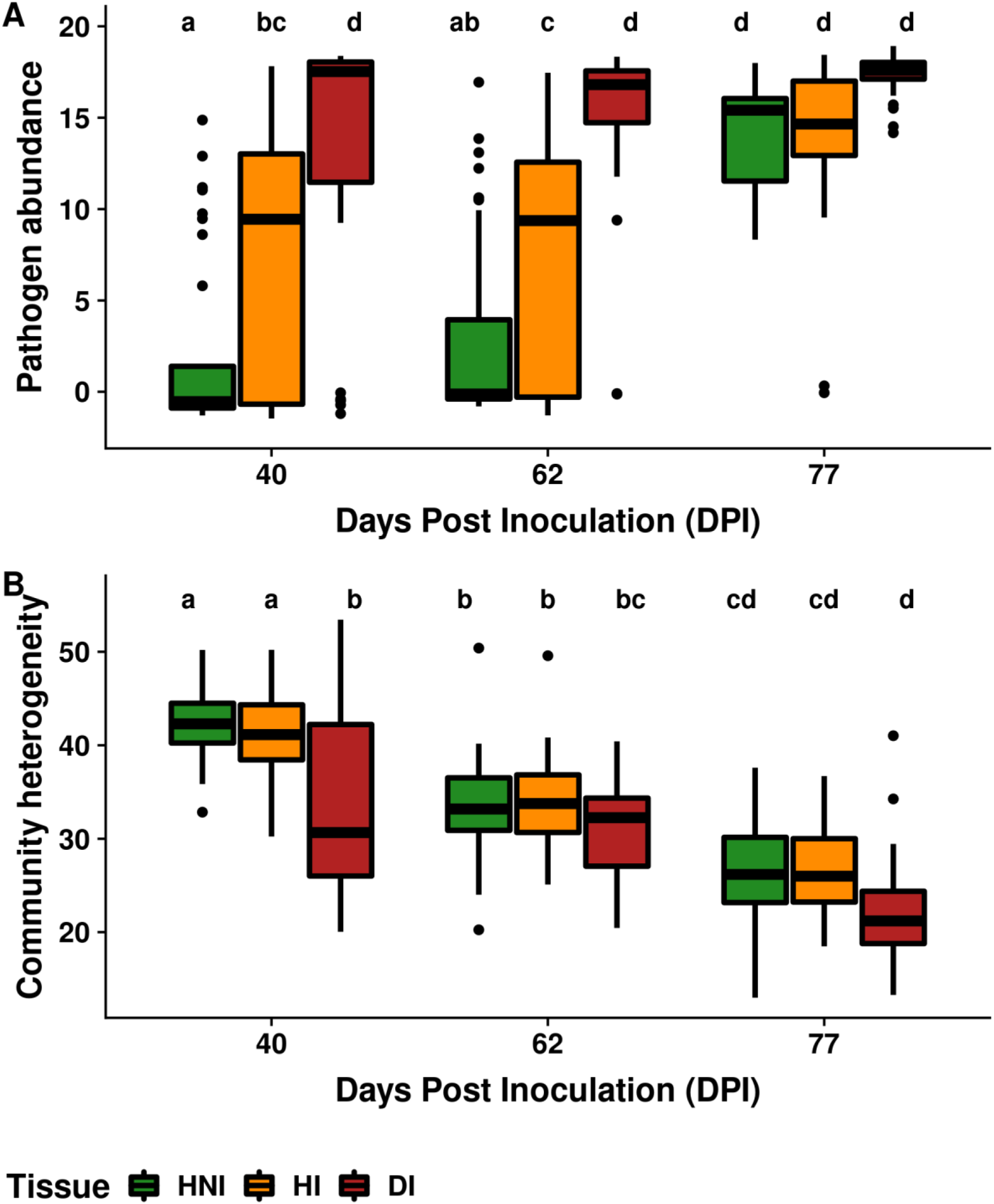
**Variations in (A) pathogen relative abundance and (B) fungal community heterogeneity among tissue types over time.** Pathogen abundance is the number of sequences assigned to *E. necator* (clr-transformed), while community heterogeneity represents the compositional similarity between each sample and the centroid of its group. HNI corresponds to the visually healthy leaf blade of non-infected leaves, while HI and DI corresponds to the visually healthy leaf blade and disease spots of infected leaves, respectively. Different letters indicate significant post-hoc comparisons.

**Figure S2.**
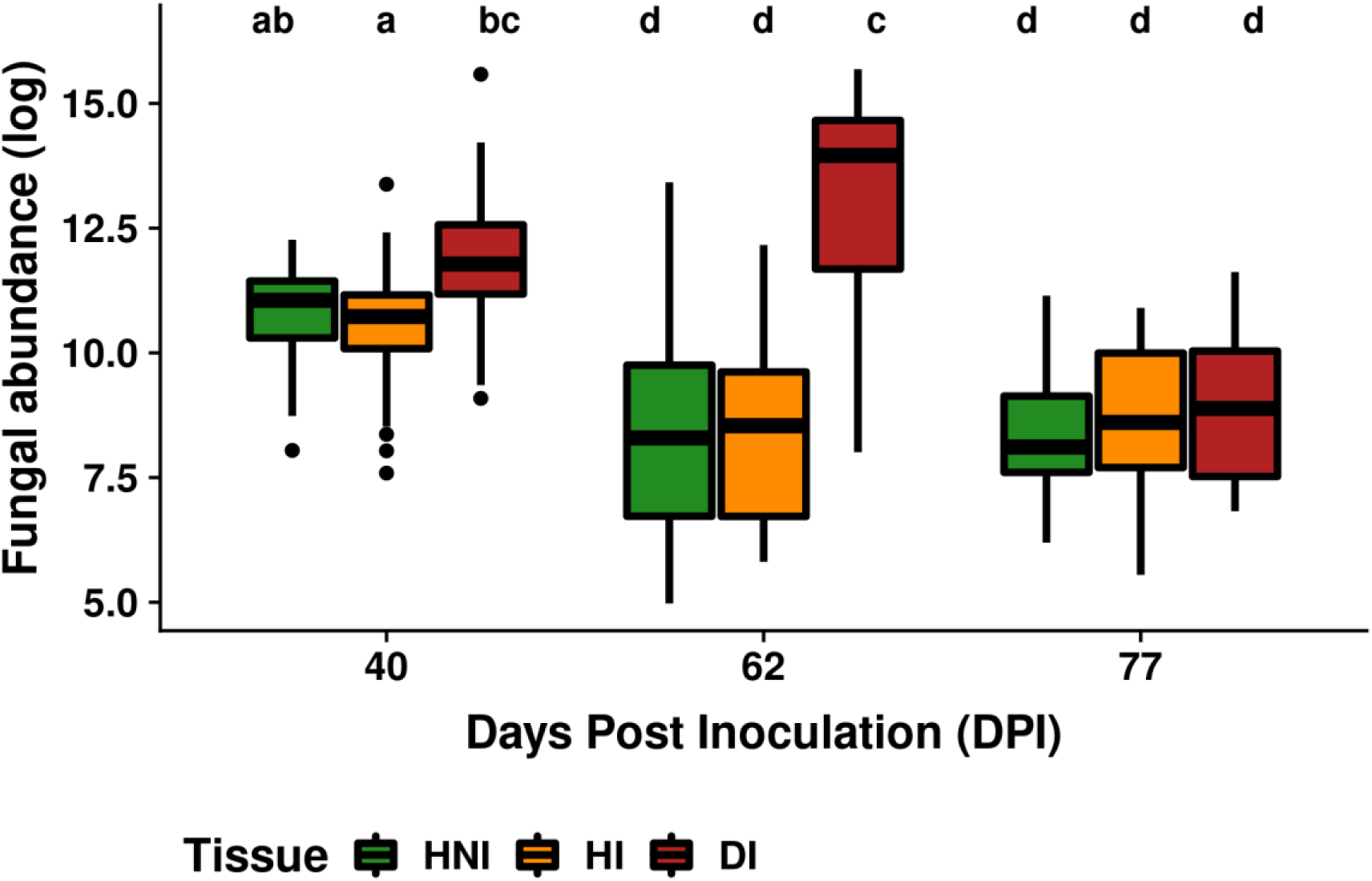
**Variations in fungal total abundance among tissue types over time.** Fungal total abundance is the number of fungal ITS1F copies (log-transformed) estimated by ddPCR. HNI corresponds to the visually healthy leaf blade of non-infected leaves, while HI and DI corresponds to the visually healthy leaf blade and disease spots of infected leaves, respectively. Different letters indicate significant post-hoc comparisons.

**Figure S3.**
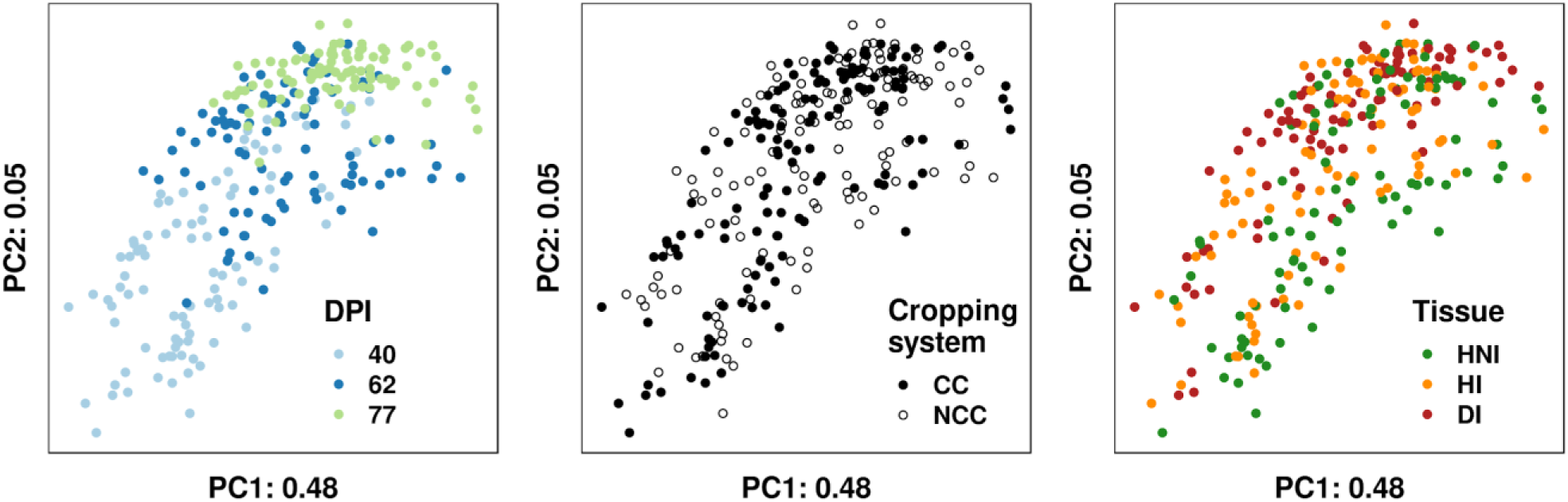
**Variations in fungal community composition of grapevine leaves between dates, cropping systems and tissue type.** Dissimilarities in composition among foliar samples were estimated with the Aitchison distance and represented with a principal coordinate analysis. Percentage variance explained by the two first axes (PC1 and PC2) are indicated. Foliar samples were collected after 40, 62 and 77 days post-inoculation (DPI), in experimental units with cover crop (CC) or without cover crop (NCC), and in three tissue types. HNI corresponds to the visually healthy leaf blade of non-infected leaves, while HI and DI corresponds to the visually healthy leaf blade and disease spots of infected leaves, respectively (Figure 1). Date and cropping system had significant effects on community composition but not tissue type (Table 1C).

**Figure S4.**
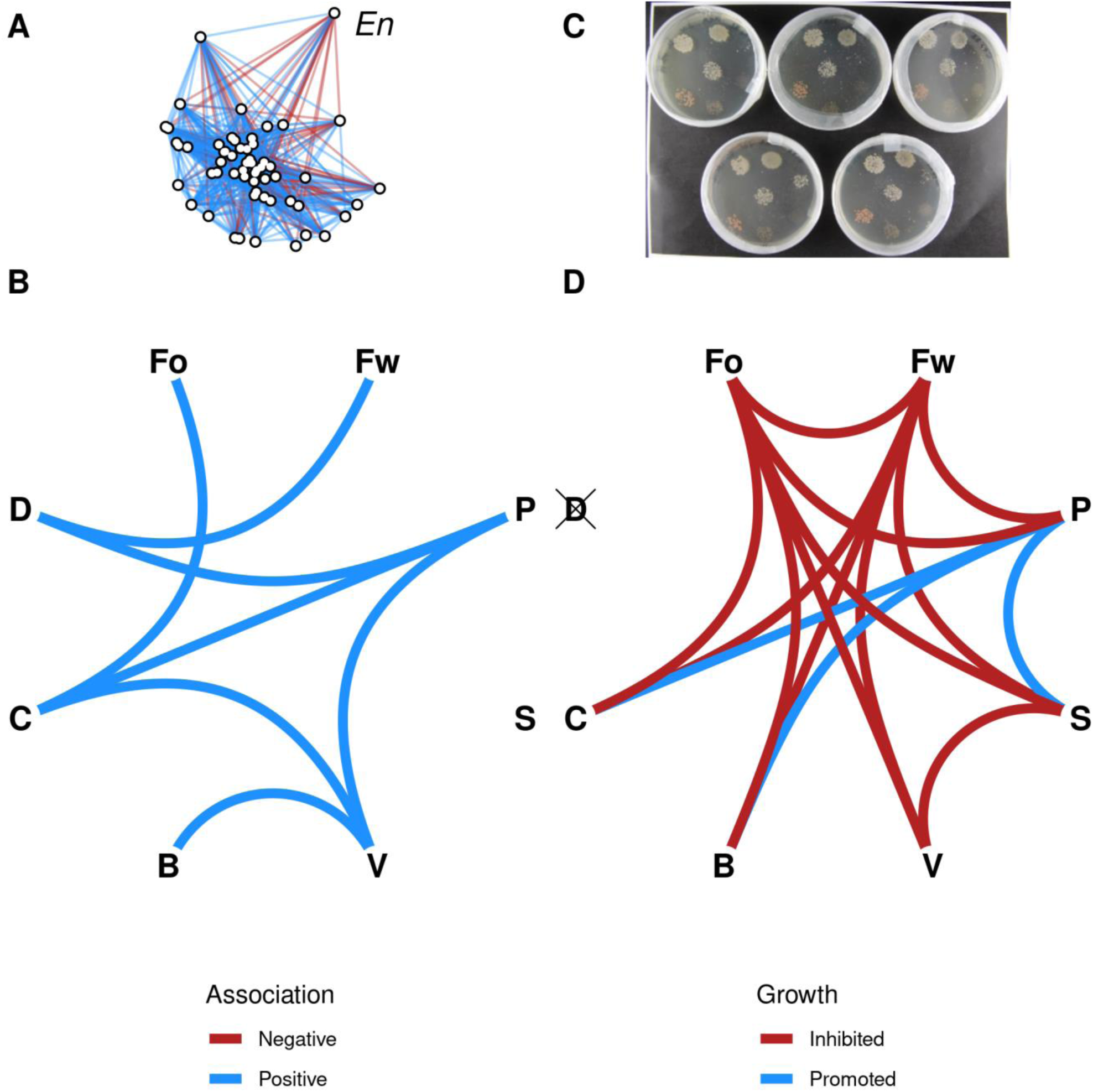
**Association network versus *in vitro* interaction network among eight yeast species.** The yeast species selected for the comparison were *Buckleyzyma aurantiaca* (B), *Cystofilobasidium macerans* (C), *Dioszegia hungarica* (D), *Filobasidium oeirense* (Fo), *Filobasidium wieringae* (Fw), *Udeniomyces pyricola* (P), *Sporobolomyces roseus* (S) and *Vishniacozyma victoriae* (V). (A) Association network of fungal species on grapevine leaves inferred with PLN from metabarcoding data and environmental covariates. The node corresponding to the pathogen *Erysiphe necator* is labelled *En*. (B) Subset of the association network involving the eight selected yeast species. This subset consisted in only positive associations (blue links). (C) Picture of the spot-on-lawn experiment used to evaluate pairwise interactions among the yeast species. A single yeast species was seeded in each Petri dish and further inoculated by spots of 8 yeast species to evaluate interactions. (D) *In vitro* interaction network among the yeast species, formed of growth-promoting (blue links) and growth-inhibiting (red links) interactions. The yeast D did not grow in the conditions of the spot-on-lawn experiment and was not included in the interaction network.

**Figure S5.**
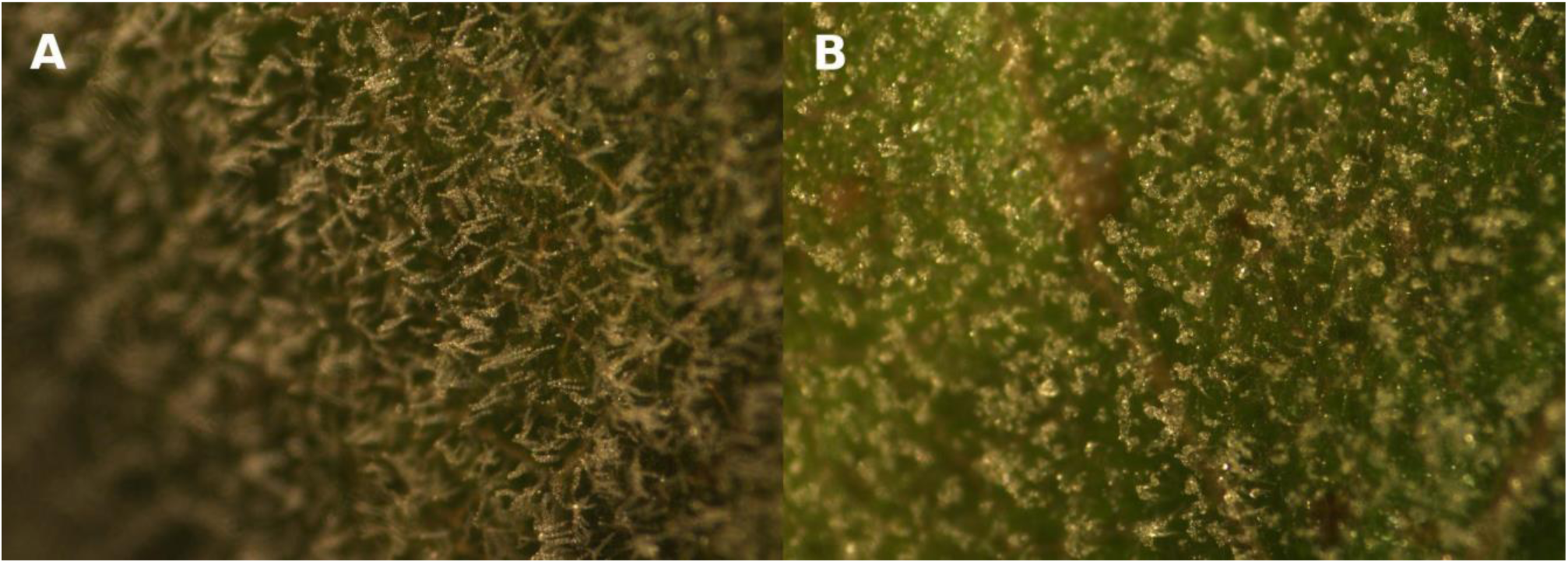
**Pictures of *E. necator* conidia in the *E. necator*-yeast strains confrontation test.** Unaltered conidial chains on control foliar discs treated with sterile distilled water (A), collapsed conidia on foliar discs treated with *B. aurantiaca* in the preventive assay (B).

**Figure S6.**
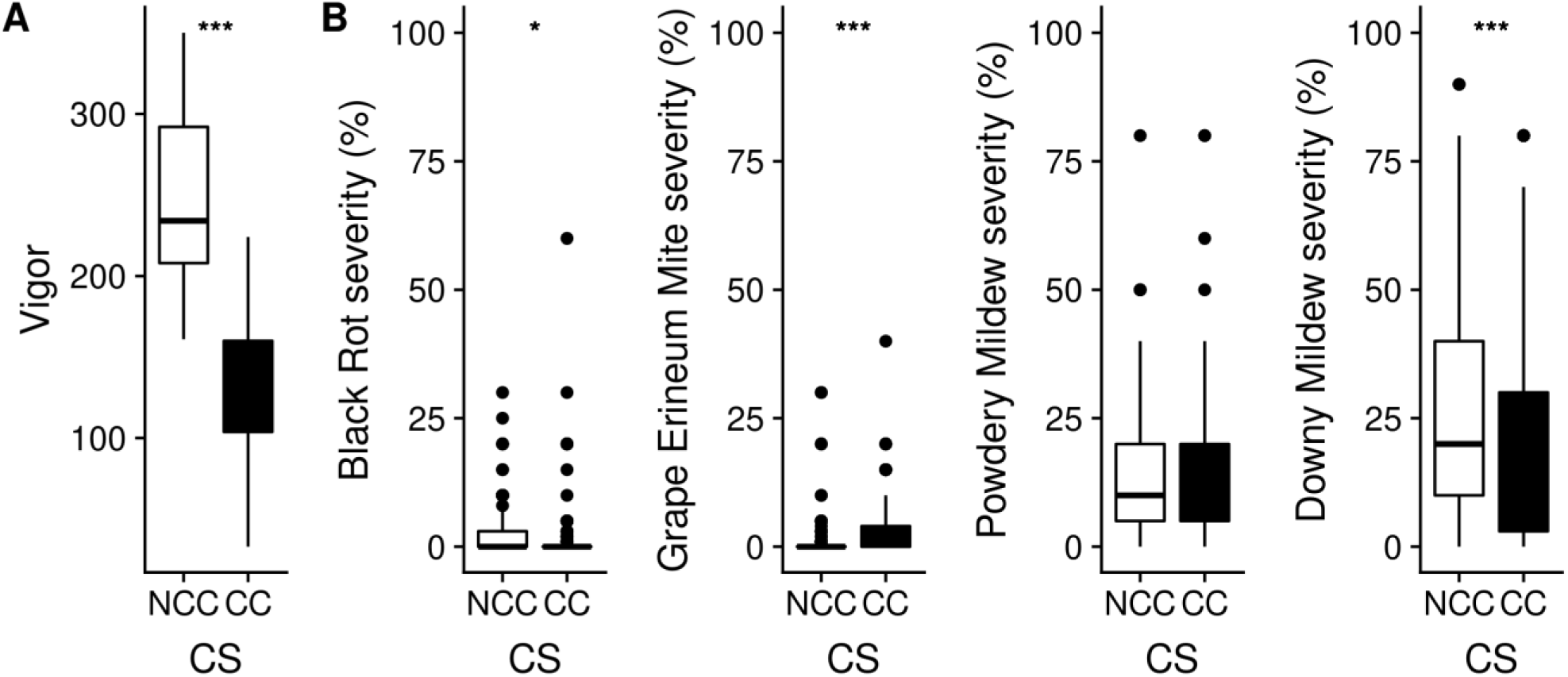
**Grapevine canopy vigor and disease severity in two cropping systems (CS), with cover crop (CC) or without cover crop (NCC).** (A) Vigor was estimated on ∼20 grapevines per cropping system. (B) Disease severity was monitored on 468 leaves randomly distributed among the two cropping systems (* p<0.05; **p<0.01; ***p<0.001).

**Table S1.** **List of the species significantly associated with *Erysiphe necator*.** Statistical associations are described with the value of the partial correlation provided by PLN and with their stability, which is the fraction of bootstrap subsamples that contained this association. The detection of the species in other microbial environments (ME) such as old leaves (OL), ground cover (GC) or bark (BK) is indicated. Significant increase in abundance of the species in a cropping system (CS) is indicated in the last column. Nine species were favored by cover cropping (CC) but no no species was favored by weed removal (NCC).

| Species | Correlation | Stability | Other ME | CS |
| --- | --- | --- | --- | --- |
| *Dioszegia butyracea* | -0.087 | 0.96 | OL / GC | CC |
| *Cladosporium ramotenellum* | -0.074 | 0.94 | OL / GC / BK |  |
| *Sawadaea bicornis* | -0.071 | 0.96 | OL |  |
| *Taphrina carpini* | -0.059 | 0.96 | OL / GC / BK | CC |
| *Stereum hirsutum* | -0.043 | 0.84 |  |  |
| *Buckleyzyma aurantiaca* | -0.035 | 0.82 | OL / GC / BK | CC |
| *Erythrobasidium hasegawianum* | -0.03 | 0.76 | OL / GC / BK | CC |
| *Taphrina deformans* | -0.02 | 0.56 | OL / GC | CC |
| *Erysiphe euonymicola* | -0.015 | 0.58 |  | CC |
| *Fomes fomentarius* | -0.011 | 0.6 | BK |  |
| *Bullera crocea* | -0.011 | 0.68 | OL / GC / BK | CC |
| *Filobasidium wieringae* | -0.01 | 0.52 | OL / GC / BK |  |
| *Neodevriesia capensis* | -0.01 | 0.38 | OL / GC / BK | CC |
| *Vishniacozyma victoriae* | -0.005 | 0.62 | OL / GC / BK | CC |
| *Itersonilia perplexans* | -0.003 | 0.38 |  |  |
| *Ascochyta manawaorae* | 0.011 | 0.52 | OL / GC / BK |  |
| *Taphrina inositophila* | 0.017 | 0.68 | OL / GC / BK |  |
| *Vishniacozyma dimennae* | 0.02 | 0.64 | OL / GC / BK |  |
| *Botrytis caroliniana* | 0.035 | 0.86 | OL / GC / BK |  |
| *Apiotrichum domesticum* | 0.038 | 0.8 |  |  |
| *Filobasidium stepposum* | 0.055 | 1 |  |  |
| *Bulleromyces albus* | 0.149 | 1 | OL / GC / BK |  |
| *Pseudopithomyces chartarum* | 0.158 | 1 | OL / GC / BK |  |

**Table S2.** **List of articles recovered by the text-mining approach from the Scopus database**. These articles had both *Erysiphe necator* and the species name of a putative antagonist in the abstract, title, keywords or references. The search included species synonyms through a custom script (Methods S1).

| Pathogen | Putative antagonist searched for | Result |
| --- | --- | --- |
| *Erysiphe necator* | *Erythrobasidium hasegawianum* | Mueller, G. M., Bills, G. F., & Foster, M. S. (Eds.). (2004). *Biodiversity of fungi: inventory and monitoring methods*. Amsterdam ; Boston: Elsevier. |
| *Erysiphe necator* | *Fomes fomentarius* | Tabata, J., De Moraes, C. M., & Mescher, M. C. (2011). Olfactory Cues from Plants Infected by Powdery Mildew Guide Foraging by a Mycophagous Ladybird Beetle. *PLoS ONE*, 6(8), e23799. doi: [10.1371/journal.pone.0023799](https://doi.org/10.1371/journal.pone.0023799) |
| *Erysiphe necator* | *Sawadaea bicornis* | Kiss, L. (1998). Natural occurrence of *Ampelomyces* intracellular mycoparasites in mycelia of powdery mildew fungi. *New Phytologist*, 140(4), 709–714. doi: [10.1046/j.1469-8137.1998.00316.x](https://doi.org/10.1046/j.1469-8137.1998.00316.x) |
| *Erysiphe necator* | *Taphrina deformans* | Chang, H.-X., Noel, Z. A., Sang, H., & Chilvers, M. I. (2018). Annotation resource of tandem repeat-containing secretory proteins in sixty fungi. *Fungal Genetics and Biology*, 119, 7–19. doi: [10.1016/j.fgb.2018.07.004](https://doi.org/10.1016/j.fgb.2018.07.004) |
| *Erysiphe necator* | *Taphrina deformans* | Kües, U., Khonsuntia, W., & Subba, S. (2018). Complex fungi. *Fungal Biology Reviews*, 32(4), 205–218. doi: [10.1016/j.fbr.2018.08.001](https://doi.org/10.1016/j.fbr.2018.08.001) |
| *Erysiphe necator* | *Taphrina deformans* | Caubel, J., Launay, M., Lannou, C., & Brisson, N. (2012). Generic response functions to simulate climate-based processes in models for the development of airborne fungal crop pathogens. *Ecological Modelling*, 242, 92–104. doi: [10.1016/j.ecolmodel.2012.05.012](https://doi.org/10.1016/j.ecolmodel.2012.05.012) |
| *Erysiphe necator* | *Taphrina deformans* | Weete, J. D., Abril, M., & Blackwell, M. (2010). Phylogenetic Distribution of Fungal Sterols. *PLoS ONE*, 5(5), e10899. doi: [10.1371/journal.pone.0010899](https://doi.org/10.1371/journal.pone.0010899) |
| *Erysiphe necator* | *Taphrina deformans* | Mysyakina, I. S., & Funtikova, N. S. (2007). The role of sterols in morphogenetic processes and dimorphism in fungi. *Microbiology*, 76(1), 1–13. doi: [10.1134/S0026261707010018](https://doi.org/10.1134/S0026261707010018) |
| *Erysiphe necator* | *Taphrina deformans* | Hernandez, A., Cooke, D. T., Lewis, M., & Clarkson, D. T. (1997). Fungicides and sterol-deficient mutants of *Ustilago maydis*: plasma membrane physico-chemical characteristics do not explain growth inhibition. *Microbiology*, 143(10), 3165–3174. doi: [10.1099/00221287-143-10-3165](https://doi.org/10.1099/00221287-143-10-3165) |

**Table S3.** **Effect of preventive or curative treatments with yeast strains on the conidia of *E. necator* in controlled conditions.** The number of foliar discs (out of 8) with altered conidia were counted for each treatment. Bold entries indicate significant differences in the number of altered conidia relative to the negative control using Fisher exact test. No conidia were observed with the positive control (benzothiadiazole) which was thus not included.

|  | Preventive treatment |  |  |  | Curative treatment |  |  |  |
| --- | --- | --- | --- | --- | --- | --- | --- | --- |
|  | Pellet |  | Supernatant |  | Pellet |  | Supernatant |  |
| Yeast strain | Disks with altered conidia | p-value | Disks with altered conidia | p-value | Disks with altered conidia | p-value | Disks with altered conidia | p-value |
| *V. victoriae* | 4 | 0.141 | 4 | 0.141 | 1 | 0.7667 | 1 | 0.7667 |
| *B. aurantiaca* | 3 | 0.2846 | 5 | 0.0594 | 5 | 0.0594 | **8** | **0.0007** |
| *D. hungarica* | 1 | 0.7667 | 1 | 0.7667 | 4 | 0.141 | 0 | 1 |
| *U. pyricola* | 1 | 0.7667 | 0 | 1 | 5 | 0.0594 | 0 | 1 |
| *S. roseus* | 1 | 0.7667 | 1 | 0.7667 | 5 | 0.0594 | 1 | 0.7667 |
| *C. macerans* | 0 | 1 | 2 | 0.5 | 3 | 0.2846 | 0 | 1 |
| *F. oeirense* | 0 | 1 | 5 | 0.0594 | 3 | 0.2846 | 0 | 1 |
| *F. wieringae* | 0 | 1 | 0 | 1 | 3 | 0.2846 | 1 | 0.7667 |
| Negative control | 1 |  | 1 |  | 1 |  | 1 |  |

**Table S4.** **List of the species differentially abundant between cropping systems (CS)**, according to ALDEX2 analysis. The table indicates the most favorable cropping system for each species (with cover crop (CC) or without cover crop (NCC)), the effect size and the Benjamini-Hochberg corrected p-value.

| Species | CS | Effect size | p-value |
| --- | --- | --- | --- |
| *Erysiphe necator* | NCC | 0.45 | 2.80E-07 |
| *Udeniomyces pyricola* | CC | -0.17 | 6.90E-02 |
| *Taphrina caerulescens* | CC | -0.17 | 7.10E-02 |
| *Blumeria graminis* | CC | -0.18 | 6.10E-02 |
| *Symmetrospora gracilis* | CC | -0.21 | 4.00E-02 |
| *Filobasidium oeirense* | CC | -0.22 | 9.30E-02 |
| *Articulospora proliferata* | CC | -0.25 | 2.60E-02 |
| *Gelidatrema spencermartinsiae* | CC | -0.26 | 3.10E-02 |
| *Dioszegia butyracea* | CC | -0.26 | 3.50E-02 |
| *Erythrobasidium hasegawianum* | CC | -0.26 | 1.30E-02 |
| *Limonomyces culmigenus* | CC | -0.27 | 1.60E-02 |
| *Vishniacozyma victoriae* | CC | -0.3 | 4.90E-04 |
| *Dioszegia hungarica* | CC | -0.3 | 4.80E-04 |
| *Neodevriesia capensis* | CC | -0.3 | 3.00E-02 |
| *Taphrina deformans* | CC | -0.31 | 1.60E-03 |
| *Bullera crocea* | CC | -0.32 | 6.10E-03 |
| *Erysiphe euonymicola* | CC | -0.39 | 1.70E-07 |
| *Buckleyzyma aurantiaca* | CC | -0.46 | 2.30E-05 |
| *Curvibasidium cygneicollum* | CC | -0.46 | 5.60E-05 |
| *Mycosphaerella tassiana* | CC | -0.51 | 6.80E-08 |
| *Taphrina carpini* | CC | -0.51 | 7.20E-06 |
| *Cystofilobasidium macerans* | CC | -0.55 | 5.10E-07 |
| *Mycosphaerella punctiformis* | CC | -0.57 | 3.50E-07 |
| *Angustimassarina acerina* | CC | -0.62 | 1.70E-08 |
| *Itersonilia pannonica* | CC | -0.74 | 5.40E-12 |

**Table S5.** **Pesticides applied in the study site since 2006 according to the pest targeted.** D: Downy mildew (*Plasmopara viticola*); P: Powdery mildew (*Erysiphe necator*), E: European grapevine moth (*Lobesia botrana*) and S: Flavescence dorée vector (*Scaphoideus titanus*). Application dates (month/day) are given into brackets.

| Pesticide active compounds | Target pest | Year (month/day) |
| --- | --- | --- |
| Chlorpyrifos-ethyl | T, S | 2006 (7/3); 2009 (5/20, 7/6) |
| Copper compounds | D | 2008 (7/30) |
| Fosetyl + Folpet | D | 2008 (7/21) |
| Lambda-cyhalothrin | S | 2009 (6/16) |
| Mancozeb + Cymoxanil | D | 2006 (5/23, 6/9); 2007 (6/11); 2008 (5/28, 6/6, 6/26); 2009 (6/26); 2011 (5/19); 2012 (6/22, 7/2) |
| Quinoxifen | P | 2008 (6/6); 2011 (5/19) |
| Spiroxamin | P | 2008 (6/26, 7/21) |

## Methods S1 R code for searching fungal species names co-occurrences in the literature

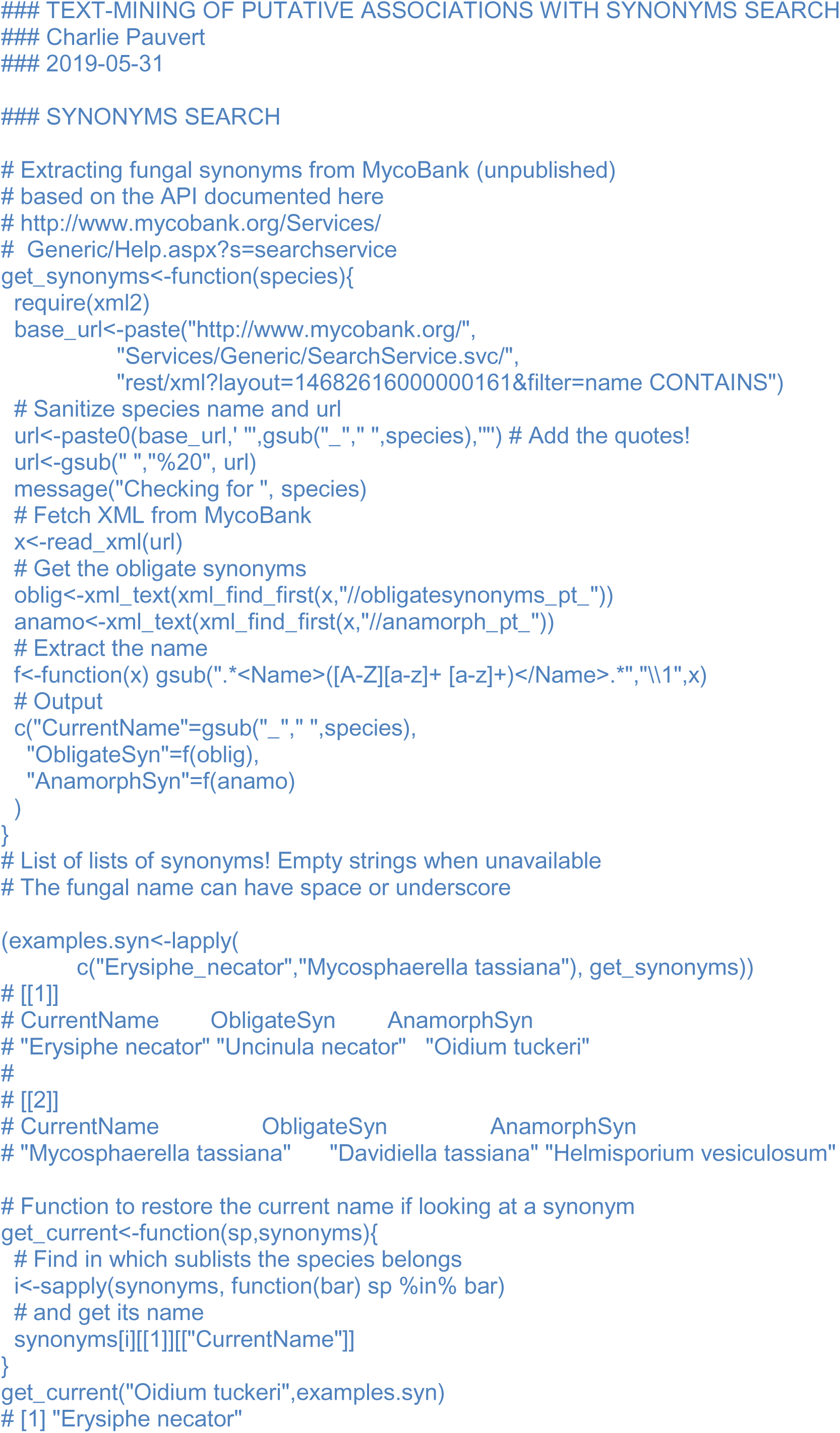
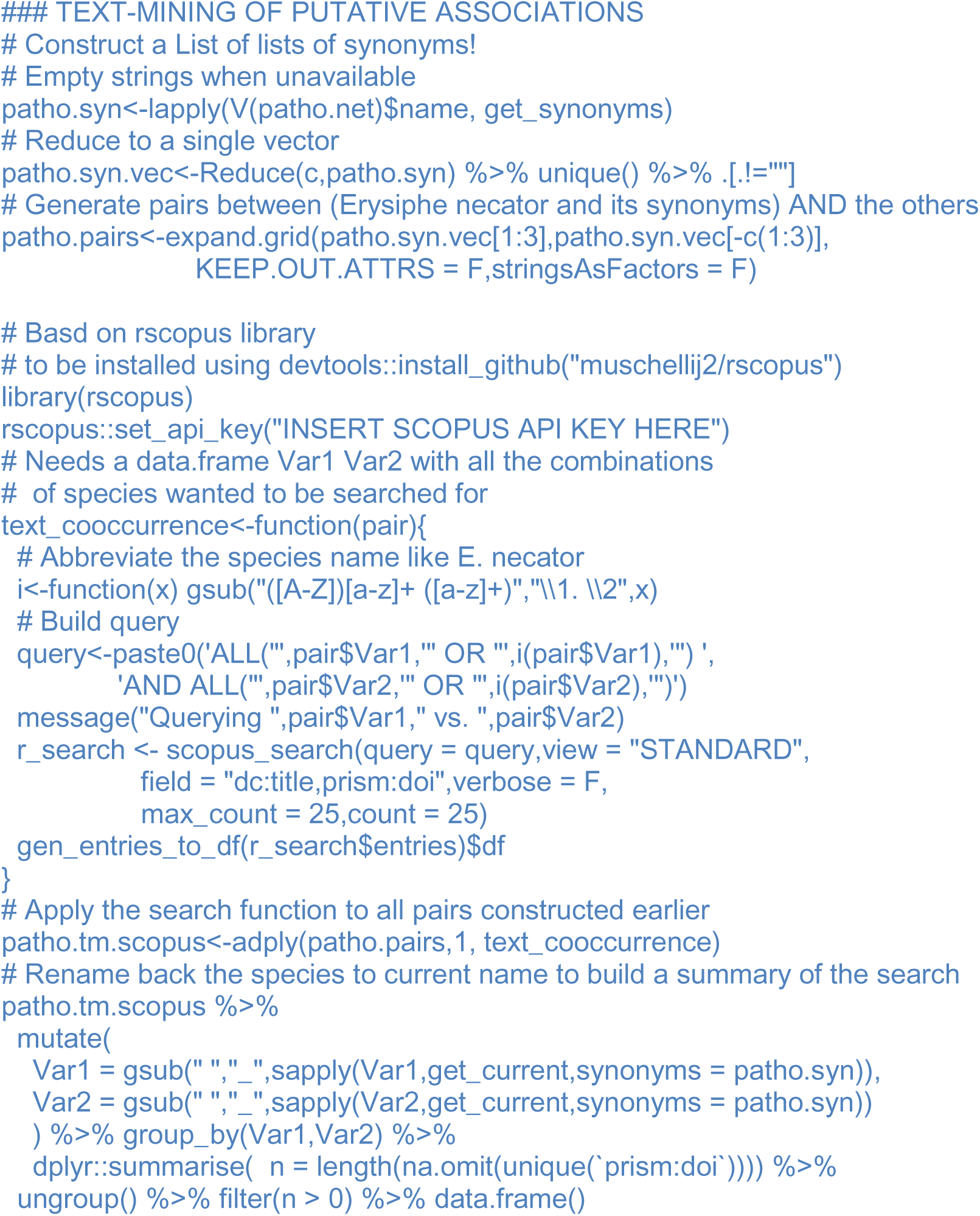

## REFERENCES

Abdelfattah, A., Malacrinò, A., Wisniewski, M., Cacciola, S.O., Schena, L. (2018) Metabarcoding: A powerful tool to investigate microbial communities and shape future plant protection strategies. Biol Control 120: 1–10.

Addis, E., Fleet, G.H., Cox, J.M., Kolak, D., Leung, T. (2001) The growth, properties and interactions of yeasts and bacteria associated with the maturation of Camembert and blue-veined cheeses. Int J Food Microbiology 69: 25–36.

Agler, M.T., Ruhe, J., Kroll, S., Morhenn, C., Kim, S-T., Weigel, D., Kemen, E.M. (2016) Microbial Hub Taxa Link Host and Abiotic Factors to Plant Microbiome Variation. PLOS Biol 14: e1002352.

Anderson, M.J. (2006) Distance-Based Tests for Homogeneity of Multivariate Dispersions. Biometrics 62: 245–253.

Anderson, M.J. (2001) A new method for non-parametric multivariate analysis of variance. Austral Ecol 26: 32–46.

Armijo, G., Schlechter, R., Agurto, M., Muñoz, D., Nuñez, C., Arce-Johnson, P. (2016) Grapevine Pathogenic Microorganisms: Understanding Infection Strategies and Host Response Scenarios. Front Plant Sci 7: 382.

Arnold, A.E., Mejía, L.C., Kyllo, D., Rojas, E.I., Maynard, Z., Robbins, N., Herre, E.A. (2003) Fungal endophytes limit pathogen damage in a tropical tree. PNAS 100: 15649–15654.

Bass, D., Stentiford, G.D., Wang, H-C., Koskella, B., Tyler, C.R. (2019) The Pathobiome in Animal and Plant Diseases. Trends Ecol Evol 34: 996–1008.

Berry, D., Widder, S. (2014) Deciphering microbial interactions and detecting keystone species with co-occurrence networks. Front Microbiol 5: 219.

Biswas, S., Mcdonald, M., Lundberg, D.S., Dangl, J.L., Jojic, V. (2016) Learning microbial interaction networks from metagenomic count data. J Comput Biol 23: 526–535.

Brader, G., Compant, S., Vescio, K., Mitter, B., Trognitz, F., Ma, L-J., Sessitsch, A. (2017) Ecology and Genomic Insights on Plant-Pathogenic and -Nonpathogenic Endophytes. Ann Rev Phytopathol 55: 61–83.

Burgaud, G., Arzur, D., Sampaio, J.P., Barbier, G. (2011) *Candida oceani sp. nov*., a novel yeast isolated from a Mid-Atlantic Ridge hydrothermal vent (−2300 meters). Antonie van Leeuwenhoek 100: 75–82.

Burgaud, G., Coton, M., Jacques, N., Debaets, S., Maciel, N.O.P., Rosa, C.A., et al. (2016) Yamadazyma barbieri f.a. sp. nov., an ascomycetous anamorphic yeast isolated from a Mid-Atlantic Ridge hydrothermal site (−2300 m) and marine coastal waters. Int J Syst Evol Micr 66: 3600–3606.

Callahan, B.J., McMurdie, P.J., Holmes, S.P. (2017) Exact sequence variants should replace operational taxonomic units in marker-gene data analysis. ISME J 11: 2639–2643.

Callahan, B.J., McMurdie, P.J., Rosen, M.J., Han, A.W., Johnson, A.J.A., Holmes, S.P. (2016) DADA2: High-resolution sample inference from Illumina amplicon data. Nat Methods 13: 581–583.

Calonnec, A., Cartolaro, P., Chadœuf, J. (2009) Highlighting Features of Spatiotemporal Spread of Powdery Mildew Epidemics in the Vineyard Using Statistical Modeling on Field Experimental Data. Phytopathology 99: 411–422.

Calonnec, A., Cartolaro, P., Delière, L., Chadœuf, J. (2006) Powdery mildew on grapevine: the date of primary contamination affects disease development on leaves and damage on grape. Bull OILB/SROP 29: 67–73.

Calonnec, A., Jolivet, J., Vivin, P., Schnee, S. (2018) Pathogenicity Traits Correlate With the Susceptible *Vitis vinifera* Leaf Physiology Transition in the Biotroph Fungus *Erysiphe necator*: An Adaptation to Plant Ontogenic Resistance. Front Plant Sci 9: 1808.

Cartolaro, P., Steva, H. (1990) Control of powdery mildew infection in grapevine in laboratory. Phytoma 419: 37–40.

Chen, J., Chen, Z. (2008) Extended Bayesian information criteria for model selection with large model spaces. Biometrika 95: 759–771.

Chiquet, J., Mariadassou, M., Robin, S. (2019) Variational inference for sparse network reconstruction from count data. PMLR 97: 1162–1171.

Chiquet, J., Mariadassou, M., Robin, S. (2018) Variational inference for probabilistic Poisson PCA. Ann Appl Stat 12: 2674–2698.

Cougoul, A., Bailly, X., Vourc’h, G., Gasqui, P. (2019) Rarity of microbial species: In search of reliable associations. PLOS ONE 14: e0200458.

Crane, S. L., van Dorst, J., Hose, G. C., King, C. K., & Ferrari, B. C. (2018). Microfluidic qPCR enables high throughput quantification of microbial functional genes but requires strict curation of primers. Front Env Sci 6: 145.

Das, P., Ji, B., Kovatcheva-Datchary, P., Bäckhed, F., Nielsen, J. (2018) In vitro co-cultures of human gut bacterial species as predicted from co-occurrence network analysis. PLOS ONE 13: e0195161.

Dannemiller, K.C., Lang-Yona, N., Yamamoto, N., Rudich, Y., Peccia, J. (2014) Combining real-time PCR and next-generation DNA sequencing to provide quantitative comparisons of fungal aerosol populations. Atmosph Environ 84: 113–121.

de Melo, E.A., de Lima, R.M.R., Laranjeira, D., dos Santos, L.A., de Omena, G.L., Barbosa de Souza, E. (2015) Efficacy of Yeast in the Biocontrol of Bacterial Fruit Blotch in Melon Plants. Trop Plant Pathol 40: 56–64.

Derocles, S.A.P., Bohan, D.A., Dumbrell, A.J., Kitson, J.J.N., Massol, F., Pauvert, C., et al. (2018) Chapter One - Biomonitoring for the 21st Century: Integrating Next-Generation Sequencing Into Ecological Network Analysis, in: Bohan, D.A., Dumbrell, A.J., Woodward, G., Jackson, M. (Eds.), Advances in Ecological Research, Next Generation Biomonitoring: Part 1. Academic Press, pp. 1–62.

Dufour, M.C., Lambert, C., Bouscaut, J., Mérillon, J.M., Corio-Costet, M.F. (2013) Benzothiadiazole-primed defence responses and enhanced differential expression of defence genes in *Vitis vinifera* infected with biotrophic pathogens *Erysiphe necator* and *Plasmopara viticola:* Elicitation and grapevine responses to mildews. Plant Pathol 62: 370–382.

Durán, P., Thiergart, T., Garrido-Oter, R., Agler, M., Kemen, E., Schulze-Lefert, P., Hacquard, S. (2018) Microbial Interkingdom Interactions in Roots Promote Arabidopsis Survival. Cell 175: 973–983.e14.

Faust, K., Raes, J. (2012) Microbial interactions: from networks to models. Nat Rev Microbiol 10: 538–550.

Fernandes, A.D., Reid, J.N., Macklaim, J.M., McMurrough, T.A., Edgell, D.R., Gloor, G.B. (2014) Unifying the analysis of high-throughput sequencing datasets: characterizing RNA-seq, 16S rRNA gene sequencing and selective growth experiments by compositional data analysis. Microbiome 2: 15.

Fierer, N., Jackson, J.A., Vilgalys, R., Jackson, R.B. (2005) Assessment of Soil Microbial Community Structure by Use of Taxon-Specific Quantitative PCR Assays. Appl Environ Microb 71: 4117–4120.

Fort, T., Pauvert, C., Zanne, A.E., Ovaskainen, O., Caignard, T., Barret, M., et al. (2019) Maternal effects and environmental filtering shape seed fungal communities in oak trees. bioRxiv 691121.

Fort, T., Robin, C., Capdevielle, X., Delière, L., Vacher, C. (2016) Foliar fungal communities strongly differ between habitat patches in a landscape mosaic. PeerJ 4: e2656.

Friedman, J., Alm, E.J. (2012) Inferring correlation networks from genomic survey data. PLoS Comput Biol 8: e1002687.

Gadoury, D.M., Cadle-Davidson, L., Wilcox, W.F., Dry, I.B., Seem, R.C., Milgroom, M.G (2012) Grapevine powdery mildew (*Erysiphe necator*): a fascinating system for the study of the biology, ecology and epidemiology of an obligate biotroph. Mol Plant Pathol 13: 1–16.

Galan, M., Razzauti, M., Bard, E., Bernard, M., Brouat, C., Charbonnel, N., et al. (2016) 16S rRNA Amplicon Sequencing for Epidemiological Surveys of Bacteria in Wildlife. mSystems 1: e00032–16.

Gardes, M., Bruns, T.D. (1993) ITS primers with enhanced specificity for basidiomycetes - application to the identification of mycorrhizae and rusts. Mol Ecol 2: 113–118.

Gloor, G.B., Macklaim, J.M., Pawlowsky-Glahn, V., Egozcue, J.J. (2017) Microbiome Datasets Are Compositional: And This Is Not Optional. Front Microbiol 8: 2224.

Gobbi, A., Kyrkou, I., Filippi, E., Ellegaard-Jensen, L., Hansen, L. H. (2020) Seasonal epiphytic microbial dynamics on grapevine leaves under biocontrol and copper fungicide treatments. Sci Rep 10: 1–13.

Hacquard, S., Spaepen, S., Garrido-Oter, R., Schulze-Lefert, P. (2017) Interplay Between Innate Immunity and the Plant Microbiota. Ann Rev Phytopathol 55: 565–589.

Hartman, K., van der Heijden, M.G.A., Wittwer, R.A., Banerjee, S., Walser, J-C., Schlaeppi, K. (2018) Cropping practices manipulate abundance patterns of root and soil microbiome members paving the way to smart farming. Microbiome 6: 14.

Hassani, M.A., Durán, P., Hacquard, S. (2018) Microbial interactions within the plant holobiont. Microbiome 6: 58.

Heard, G.M., Fleet, G.H. (1987) Occurrence and Growth of Killer Yeasts during Wine Fermentation. Appl Environ Microbiol 53: 2171–2174.

Hindson, B.J., Ness, K.D., Masquelier, D.A., Belgrader, P., Heredia, N.J., Makarewicz, A.J., et al. (2011) High-Throughput Droplet Digital PCR System for Absolute Quantitation of DNA Copy Number. Anal Chem 83: 8604– 8610.

Hirano, H., Takemoto, K. (2019) Difficulty in inferring microbial community structure based on co-occurrence network approaches. BMC Bioinformatics 20: 329.

Jacobs, B.K.M., Goetghebeur, E., Vandesompele, J., De Ganck, A., Nijs, N., Beckers, A., et al. (2017) Model-Based Classification for Digital PCR: Your Umbrella for Rain. Anal Chem 89: 4461–4467.

Jakuschkin, B., Fievet, V., Schwaller, L., Fort, T., Robin, C., Vacher, C. (2016) Deciphering the Pathobiome: Intra- and Interkingdom Interactions Involving the Pathogen *Erysiphe alphitoides*. Microb Ecol 72: 870–880.

Karasov, T.L., Neumann, M., Duque-Jaramillo, A., Kersten, S., Bezrukov, I., Schröppel, B., et al. (2019) The relationship between microbial biomass and disease in the Arabidopsis thaliana phyllosphere. BioRxiv 828814.

Kehe, J., Kulesa, A., Ortiz, A., Ackerman, C.M., Thakku, S.G., Sellers, D., et al. (2019) Massively parallel screening of synthetic microbial communities. PNAS 116: 12804–12809.

Kemen, E. (2014) Microbe–microbe interactions determine oomycete and fungal host colonization. Curr Opin Plant Biol 20: 75–81.

Kernaghan, G., Mayerhofer, M., Griffin, A. (2017) Fungal endophytes of wild and hybrid Vitis leaves and their potential for vineyard biocontrol. Can J Microbiol 63: 583–595.

Kleyer, H., Tecon, R., Or, D. (2017) Resolving species level changes in a representative soil bacterial community using microfluidic quantitative PCR. Frontiers Microbiol 8: 2017.

Lagier, J-C., Dubourg, G., Million, M., Cadoret, F., Bilen, M., Fenollar, F., et al. (2018) Culturing the human microbiota and culturomics. Nat Rev Microbiol 16: 540–550.

Layeghifard, M., Hwang, D.M., Guttman, D.S. (2017) Disentangling Interactions in the Microbiome: A Network Perspective. Trends Microbiol 25: 217–228.

Lee, G., Lee, S-H., Kim, K.M., Ryu, C-M. (2017) Foliar application of the leaf-colonizing yeast *Pseudozyma churashimaensis* elicits systemic defense of pepper against bacterial and viral pathogens. Sci Rep 7: 39432.

Lenth, R. (2018) Emmeans: estimated marginal means, aka least-squares means.

Li, C., Lim, K.M.K., Chang, K.R., Nagarajan, N. (2016) Predicting microbial interactions through computational approaches. Methods 102: 12–19.

Lima-Mendez, G., Faust, K., Henry, N., Decelle, J., Colin, S., Carcillo, F., et al. (2015) Determinants of community structure in the global plankton interactome. Science 348: 1262073.

Liu, H., Roeder, K., Wasserman, L. (2010) Stability Approach to Regularization Selection (StARS) for High Dimensional Graphical Models, in: Proceedings of the 23rd International Conference on Neural Information Processing Systems - Volume 2, NIPS’10. Curran Associates Inc., USA, pp. 1432–1440.

Lo, C., Marculescu, R. (2017) MPLasso: Inferring microbial association networks using prior microbial knowledge. PLOS Comput Biol 13: e1005915.

Martin, M. (2011) Cutadapt removes adapter sequences from high-throughput sequencing reads. EMBnet.journal 17: 10.

Massart, S., Martinez-Medina, M., Jijakli, M. H. (2015) Biological control in the microbiome era: challenges and opportunities. Biol control 89: 98–108.

McMurdie, P.J., Holmes, S. (2013) phyloseq: An R Package for Reproducible Interactive Analysis and Graphics of Microbiome Census Data. PLOS ONE 8: e61217.

Muschelli, J. (2019) rscopus: Scopus Database “API” Interface.

Nilsson, R.H., Anslan, S., Bahram, M., Wurzbacher, C., Baldrian, P., Tedersoo, L. (2019) Mycobiome diversity: high-throughput sequencing and identification of fungi. Nature Rev Microbiol 17: 95–109.

Oksanen, J., Blanchet, F.G., Friendly, M., Kindt, R., Legendre, P., McGlinn, D., et al. (2018) vegan: Community Ecology Package.

Ovaskainen, O., Tikhonov, G., Norberg, A., Blanchet, F.G., Duan, L., Dunson, D., et al. (2017) How to make more out of community data? A conceptual framework and its implementation as models and software. Ecol Lett 20: 561– 576.

Panstruga, R., Kuhn, H. (2019) Mutual interplay between phytopathogenic powdery mildew fungi and other microorganisms. Mol Plant Pathol 20: 463–470.

Pauvert, C., Buée, M., Laval, V., Edel-Hermann, V., Fauchery, L., Gautier, A., et al. (2019) Bioinformatics matters: The accuracy of plant and soil fungal community data is highly dependent on the metabarcoding pipeline. Fungal Ecol. 41: 23–33.

Pertot, I., Caffi, T., Rossi, V., Mugnai, L., Hoffmann, C., Grando, M. S., et al. (2017) A critical review of plant protection tools for reducing pesticide use on grapevine and new perspectives for the implementation of IPM in viticulture. Crop Prot 97: 70–84.

Pinto, C., Gomes, A. C. (2016) *Vitis vinifera* microbiome: from basic research to technological development. BioControl 61: 243–256.

Polonelli, L., Morace, G. (1986) Reevaluation of the yeast killer phenomenon. J Clin Microbiol 24: 866–869.

Poudel, R., Jumpponen, A., Schlatter, D.C., Paulitz, T.C., Gardener, B.B.M., Kinkel, L.L., et al. (2016) Microbiome Networks: A Systems Framework for Identifying Candidate Microbial Assemblages for Disease Management. Phytopathology 106: 1083–1096.

Props, R., Kerckhof, F.M., Rubbens, P., De Vrieze, J., Sanabria, E. H., Waegeman, W., et al. (2017) Absolute quantification of microbial taxon abundances. ISME J 11: 584–587.

R Core Team (2018) R: A Language and Environment for Statistical Computing. R Foundation for Statistical Computing, Vienna, Austria.

Robert, V., Vu, D., Amor, A.B.H., van de Wiele, N., Brouwer, C., Jabas, B., et al. (2013) MycoBank gearing up for new horizons. IMA Fungus 4: 371.

Rognes, T., Flouri, T., Nichols, B., Quince, C., Mahé, F. (2016) VSEARCH: a versatile open source tool for metagenomics. PeerJ 4: e2584.

Röttjers, L., Faust, K. (2018) From hairballs to hypotheses–biological insights from microbial networks. FEMS Microbiol Lett 42: 761–780.

Sáenz-Romo, M.G., Veas-Bernal, A., Martínez-García, H., Ibáñez-Pascual, S., Martínez-Villar, E., Campos-Herrera, R., et al. (2019) Effects of Ground Cover Management on Insect Predators and Pests in a Mediterranean Vineyard. Insects 10: 421.

Schoch, C.L., Seifert, K.A., Huhndorf, S., Robert, V., Spouge, J.L., Levesque, C.A., et al. (2012) Nuclear ribosomal internal transcribed spacer (ITS) region as a universal DNA barcode marker for Fungi. PNAS 109: 6241–6246.

Thorsen, J., Brejnrod, A., Mortensen, M., Rasmussen, M.A., Stokholm, J., Al-Soud, W.A., et al. (2016) Large-scale benchmarking reveals false discoveries and count transformation sensitivity in 16S rRNA gene amplicon data analysis methods used in microbiome studies. Microbiome 4: 62.

Tipton, L., Müller, C.L., Kurtz, Z.D., Huang, L., Kleerup, E., Morris, A., et al. (2018) Fungi stabilize connectivity in the lung and skin microbial ecosystems. Microbiome 6: 12.

UNITE Community (2019) UNITE general FASTA release for Fungi 2.

Vacher, C., Tamaddoni-Nezhad, A., Kamenova, S., Peyrard, N., Moalic, Y., Sabbadin, R., et al. (2016) Learning Ecological Networks from Next-Generation Sequencing Data, in: Advances in Ecological Research. pp. 1–39.

Valdés-Gómez, H., Gary, C., Cartolaro, P., Lolas-Caneo, M., Calonnec, A. (2011) Powdery mildew development is positively influenced by grapevine vegetative growth induced by different soil management strategies. Crop Prot 30: 1168–1177.

Vandenkoornhuyse, P., Quaiser, A., Duhamel, M., Le Van, A., & Dufresne, A. (2015) The importance of the microbiome of the plant holobiont. New Phytol 206: 1196–1206.

Vandeputte, D., Kathagen, G., D’hoe, K., Vieira-Silva, S., Valles-Colomer, M., Sabino, J., et al. (2017) Quantitative microbiome profiling links gut community variation to microbial load. Nature 551: 507–511.

Vannier, N., Agler, M., Hacquard, S. (2019) Microbiota-mediated disease resistance in plants. PLOS Pathog 15: e1007740.

Vayssier-Taussat, M., Albina, E., Citti, C., Cosson, J-F., Jacques, M-A., Lebrun, M-H., et al. (2014) Shifting the paradigm from pathogens to pathobiome: new concepts in the light of meta-omics. Front Cell Infect Microbiol 4: 29.

Vogel, C., Bodenhausen, N., Gruissem, W., Vorholt, J.A. (2016) The Arabidopsis leaf transcriptome reveals distinct but also overlapping responses to colonization by phyllosphere commensals and pathogen infection with impact on plant health. New Phytol 212: 192–207.

Wang, Q., Garrity, G.M., Tiedje, J.M., Cole, J.R. (2007) Naïve Bayesian Classifier for Rapid Assignment of rRNA Sequences into the New Bacterial Taxonomy. Appl Environ Microbiol 73: 5261–5267.

Weiss, S., Van Treuren, W., Lozupone, C., Faust, K., Friedman, J., Deng, Y., et al. (2016) Correlation detection strategies in microbial data sets vary widely in sensitivity and precision. ISME J 10: 1669–1681.

White, T.J., Bruns, T., Lee, S., Taylor, J. (1990) Amplification and direct sequencing of fungal ribosomal RNA genes for phylogenetics. PCR Protocols: A Guide to Methods and Applications 315–322.

Zaneveld, J.R., McMinds, R., Vega Thurber R. (2017) Stress and stability: applying the Anna Karenina principle to animal microbiomes. Nature Microbiol 2: 17121.

Zarraonaindia, I., Owens, S. M., Weisenhorn, P., West, K., Hampton-Marcell, J., Lax, S., et al. (2015) The soil microbiome influences grapevine-associated microbiota. MBio 6: e02527–14.

